# Ethanolaminephosphate cytidyltransferase is essential for survival, lipid homeostasis and stress tolerance in *Leishmania major*

**DOI:** 10.1101/2023.01.10.523530

**Authors:** Somrita Basu, Mattie Pawlowic, Fong-Fu Hsu, Geoff Thomas, Kai Zhang

## Abstract

Glycerophospholipids including phosphatidylethanolamine (PE) and phosphatidylcholine (PC) are vital components of biological membranes. Trypanosomatid parasites of the genus *Leishmania* can acquire PE and PC via *de novo* synthesis and the uptake/remodeling of host lipids. In this study, we investigated the ethanolaminephosphate cytidyltransferase (EPCT) in *Leishmania major*, which is the causative agent for cutaneous leishmaniasis. EPCT is a key enzyme in the ethanolamine branch of the Kennedy pathway which is responsible for the *de novo* synthesis of PE. Our results demonstrate that *L. major* EPCT is a cytosolic protein capable of catalyzing the formation of CDP-ethanolamine from ethanolamine-phosphate and cytidine triphosphate. Genetic manipulation experiments indicate that EPCT is essential in both the promastigote and amastigote stages of *L. major* as the chromosomal null mutants cannot survive without the episomal expression of EPCT. This differs from our previous findings on the choline branch of the Kennedy pathway (responsible for PC synthesis) which is required only in promastigotes but not amastigotes. While episomal EPCT expression does not affect promastigote proliferation under normal conditions, it leads to reduced production of ethanolamine plasmalogen or plasmenylethanolamine, the dominant PE subtype in *Leishmania*. In addition, parasites with epsiomal EPCT exhibit heightened sensitivity to acidic pH and starvation stress, and significant reduction in virulence. In summary, our investigation demonstrates that proper regulation of EPCT expression is crucial for PE synthesis, stress response, and survival of *Leishmania* parasites throughout their life cycle.

**AUTHOR SUMMARY:** In nature, *Leishmania* parasites alternate between fast replicating, extracellular promastigotes in sand fly gut and slow growing, intracellular amastigotes in macrophages. Previous studies suggest that promastigotes acquire most of their lipids via *de novo* synthesis whereas amastigotes rely on the uptake and remodeling of host lipids. Here we investigated the function of ethanolaminephosphate cytidyltransferase (EPCT) which catalyzes a key step in the *de novo* synthesis of phosphatidylethanolamine (PE) in *Leishmania major*. Results showed that EPCT is indispensable for both promastigotes and amastigotes, indicating that *de novo* PE synthesis is still needed at certain capacity for the intracellular form of *Leishmania* parasites. In addition, elevated EPCT expression alters overall PE synthesis and compromises parasite’s tolerance to adverse conditions and is deleterious to the growth of intracellular amastigotes. These findings provide new insight into how *Leishmania* acquire essential phospholipids and how disturbance of lipid metabolism can impact parasite fitness.

## INTRODUCTION

Protozoan parasites of the genus *Leishmania* are transmitted through the bite of hematophagous sand flies. During their life cycle, *Leishmania* parasites alternate between flagellated, extracellular promastigotes in sand fly midgut and non-flagellated, intracellular amastigotes in mammalian macrophages. These parasites cause leishmaniasis which ranks among the top ten neglected tropical diseases with 10-12 million people infected and 350 million people at the risk of acquiring infection [1, 2]. Drugs for leishmaniasis are plagued with strong toxicity, low efficacy, and high cost [3, 4]. To develop better treatment, it is necessary to gain insights into how *Leishmania* acquire essential nutrients and proliferate in the harsh environment in the vector host and mammalian host.

To sustain growth, *Leishmania* parasites must generate abundant amounts of lipids including glycerophospholipids, sterols and sphingolipids. Phosphatidylethanolamine (PE) and phosphatidylcholine (PC) are two common classes of glycerophospholipids. Besides being major membrane constituents, PE and PC can function as precursors for several signaling molecules and metabolic intermediates including lyso-phospholipids, phosphatidic acid, diacylglycerol, and free fatty acids [5, 6]. In addition, PE contributes to the synthesis of GPI-anchored proteins in trypanosomatid parasites by providing the ethanolamine phosphate bridge that links proteins to glycan anchors [7]. PE is also involved in the formation of autophagosome during differentiation and starvation in *Leishmania major* [8, 9] and the posttranslational modification of eukaryotic elongation factor 1A in *T. brucei* [10].

For many eukaryotic cells, the majority of PE and PC are synthesized *de novo* via the Kennedy pathway (Fig. 1) [11]. In *Leishmania*, the key metabolite ethanolamine phosphate (EtN-P) is generated from of the sphingoid base metabolism or the phosphorylation of ethanolamine (EtN) by ethanolamine kinase [12](Fig. 1). EtN-P is then conjugated to cytidine triphosphate (CTP) to produce CDP-EtN and pyrophosphate by the enzyme ethanolaminephosphate cytidyltransferase (EPCT) [13]. In the EtN branch of the Kennedy pathway, through the activity of ethanolamine phosphotransferase (EPT), CDP-EtN is combined with 1-alkyl-2-acyl-glycerol to generate plasmenylethanolamine (PME) or ethanolamine plasmalogen, the dominant subtype of PE in *Leishmania* [14]. CDP-EtN can also be combined with 1,2-diacylglycerol to form 1,2-diacyl-PE (a minor subtype of PE in *Leishmania*) by choline ethanolamine phosphotransferase (CEPT) [15, 16]. A similar branch of the Kennedy pathway is responsible for the *de novo* synthesis of PC (choline → choline-phosphate → CDP-choline → PC), with CEPT catalyzing the last step of conjugating CDP-choline and diacylglycerol into PC as a dual activity enzyme [16-18]. Besides the Kennedy pathway, PE may be generated from the decarboxylation of phosphatidylserine (PS) in the mitochondria and PC may be generated from the N-methylation of PE [19-21].

**Figure 1.**
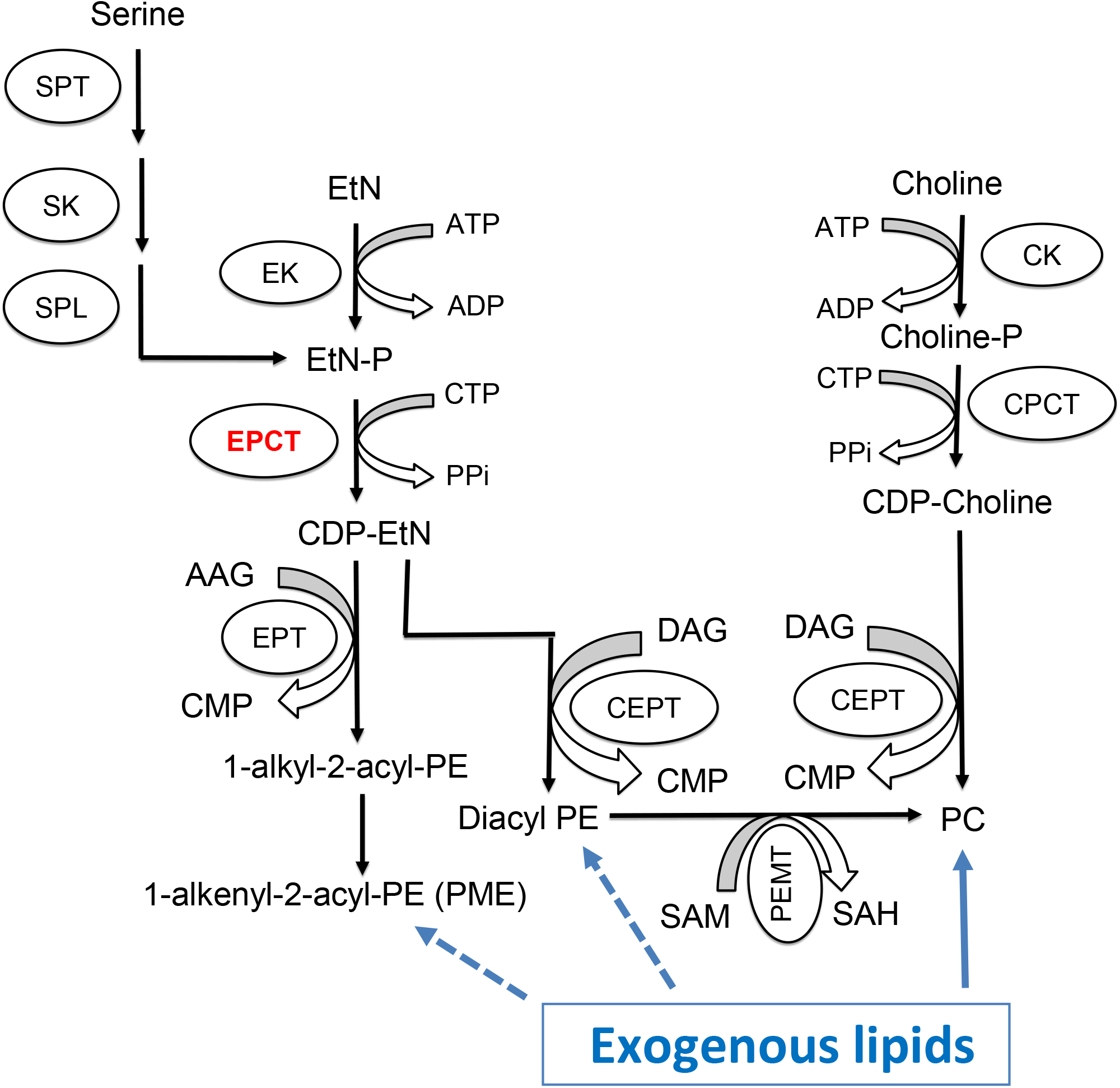
Synthesis of PE and PC in *Leishmania*. SPT: Serine palmitoyltransferase; SK: Sphingosine kinase; SPL: sphingosine-1-phosphate lyase; EK: ethanolamine kinase; EPCT: ethanolaminephosphate cytidyltransferase; EPT: ethanolamine phosphotransferase; CK: choline kinase; CPCT: cholinephosphate cytidyltransferase; CEPT: choline ethanolamine phosphotransferase; PEMT: Phosphatidylethanolamine N-methyltransferase; EtN: ethanolamine; EtN-P: ethanolamine phosphate; AAG: 1-alkyl2-acyl glycerol; DAG: 1,2-diacylglycerol; CTP: cytidine triphosphate; CDP: cytidine diphosphate; CMP: cytidine monophosphate; PPi: pyrophosphate; PME: plasmenylethanolamine; diacyl PE: 1,2-diacyl-phosphatidylethanolamine, PC: 1,2-diacyl-phosphatidylcholine; SAM: S-adenosyl-L-methionine; SAH: S-adenosyl-L-homocysteine.

In addition to biosynthesis, *Leishmania* parasites (especially the intracellular amastigotes) can take up and modify existing lipids to fulfill their needs [22-24]. For example, the enzyme CEPT which is directly responsible for producing PC and diacyl-PE (Fig. 1) is essential for *L. major* promastigotes in culture but is not required for the survival or proliferation of amastigotes in mice [18]. While diacyl-PE is a minor lipid component in *Leishmania*, PC is highly abundant in both promastigotes and amastigotes [25, 26]. These results argue that *de novo* PC synthesis is required to generate large amount of lipids to support rapid parasite replication during the promastigote stage (estimated doubling time: 6–8 hours in culture and 10–12 hours in sand fly) [27, 28]. In contrast, intracellular amastigotes can acquire enough PC through the uptake and remodeling of host lipids, which seems to fit their slow growing, metabolically quiescent state (estimated doubling time: 60 hours) [29, 30]. Consistent with these findings, sphingolipid analyses of *L. major* amastigotes revealed high levels of sphingomyelin which could not be synthesized by *Leishmania* but was plentiful in mammalian cells [31]. Conversely, the biosynthesis of parasite-specific sphingolipid, inositol phosphorylceramide (IPC), is fully dispensable for *L. major* amastigotes [12, 31, 32]. Collectively, these data suggest that as *Leishmania* transition from fast-replicating promastigotes to slow-growing amastigotes, they undergo a metabolic switch from *de novo* synthesis to scavenge/remodeling to acquire their lipids [24].

Despite of these findings, it is important to explore whether intracellular amastigotes retain some capacity for *de novo* lipid synthesis. Our previous studies on the sterol biosynthetic mutant *c14dm*^−^ suggest that is the case [33]. *L. major c14dm*^−^ mutants lack the sterol-14α-demethylase which catalyzes the removal of C-14 methyl group from sterol intermediates. Promastigotes of *c14dm*^−^ are devoid of ergostane-type sterols (which are abundant in wild type [WT] parasites) and accumulate 14-methyl sterol intermediates. However, lipid analyses of lesion-derived amastigotes demonstrate that both WT and *c14dm*^−^ amastigotes contain cholesterol (which must be taken from the host since *Leishmania* only synthesize ergostane-type sterols) as their main sterol [33]. Parasite-specific sterols, i.e., ergostane-type sterols such as ergosterol and 5-dehydroepisterol in WT amastigotes and C-14-methylated sterols in *c14dm*^−^ amastigotes are only detected at trace levels. Nonetheless, *c14dm*^−^ amastigotes show significantly attenuated virulence and reduced growth in comparison to WT and *C14DM* add-back amastigotes, suggesting that the residual amounts of endogenous sterols (which cannot be scavenged from the host) play pivotal roles in amastigotes which may constitute the basis for sterol biosynthetic inhibitors as anti-leishmaniasis drugs [33, 34].

In this study, we investigated the roles of EPCT (EC 2.7.7.14) which catalyzes the conversion of EtN-P and CTP into CDP-EtN and pyrophosphate in the Kennedy pathway (Fig. 1). Our previous study on EPT in *L. major* revealed that EPT is mostly required for the synthesis of PME but not diacyl PE or PC [14]. Disruption of EPCT, on the other hand, may have a much more profound impact on the synthesis of PME, diacyl PE and PC (which can be generated from diacyl PE via N-methylation) (Fig. 1). In mammalian cells, EPCT is considered as the rate limiting enzyme in the EtN branch of the Kennedy pathway and a key regulator of PE synthesis [35, 36]. Disruption of EPCT in mice leads to developmental defects and embryonic lethality [37, 38]. Additional studies report EPCT heterozygous mice have increased diacylglycerol and triacylglycerol in liver that resemble features of metabolic diseases, suggesting that EPCT is involved in maintaining the homeostasis of neutral lipids as well [38, 39]. Based on these findings, it is of interest to determine the roles of EPCT in *Leishmania* especially during the disease-causing intracellular stage when parasites acquire most of their lipids via scavenging. Our results indicate that EPCT is essential for the survival of both promastigotes and amastigotes in *L. major*. In addition, the expression level of EPCT is crucial for stress tolerance, lipid homeostasis and the synthesis of GPI-anchored proteins. These findings may inform the development of new anti-*Leishmania* drugs.

## RESULTS

### Identification and functional verification of *L. major* EPCT

A putative *EPCT* gene was identified in the genome of *L. major* (Tritrypdb: LmjF 32.0890) with synthetic orthologs in other *Leishmania* spp., *Trypanosoma brucei*, and *Trypanosoma cruzi*. The predicted *L. major* EPCT protein consists of 402 amino acids and bears 35-42% identity to EPCTs from *Saccharomyces cerevisiae, Plasmodium falciparum, Arabidopsis thaliana*, and *Homo sapiens* (Fig. S1).

To confirm its function, the *EPCT* open reading frame (ORF) was cloned into a pXG plasmid (a high copy number protein expression vector) [40] and introduced into *L. major* wild type parasites (WT) as WT+EPCT. When promastigote lysate from WT+EPCT was incubated with [^14^C]-EtN-P and CTP for 20 minutes at room temperature, we detected a radiolabeled product which exhibited similar mobility (retention factor) as pure CDP-EtN on thin layer chromatography (TLC), suggesting it was [^14^C]-CDP-EtN (Fig. 2A and Fig. S2A). Radiolabeled CDP-EtN was also observed when [^14^C]-EtN-P and CTP were incubated with mouse liver homogenate, but not boiled lysates (Fig. 2A and Fig. S2B). We did not detect the formation of [^14^C]-CDP-EtN using WT lysate, indicating that the basal level of EPCT activity in *L. major* promastigotes was below the level of detection with this approach (Fig. 2A). To determine the cellular localization of EPCT, a GFP-tagged EPCT was introduced into WT promastigotes as WT+GFP-EPCT. As shown in Fig. 2B-C, GFP-EPCT was detected at the predicted molecular weight of ∼73 kDa and exhibited a cytoplasmic localization. In addition, WT+GFP-EPCT cells displayed similar EPCT activity levels as WT+EPCT cells (Fig. 2A). Together, these findings argue that *L. major* EPCT is a cytosolic enzyme capable of condensing EtN-P and CTP into CDP-EtN.

**Figure 2.**
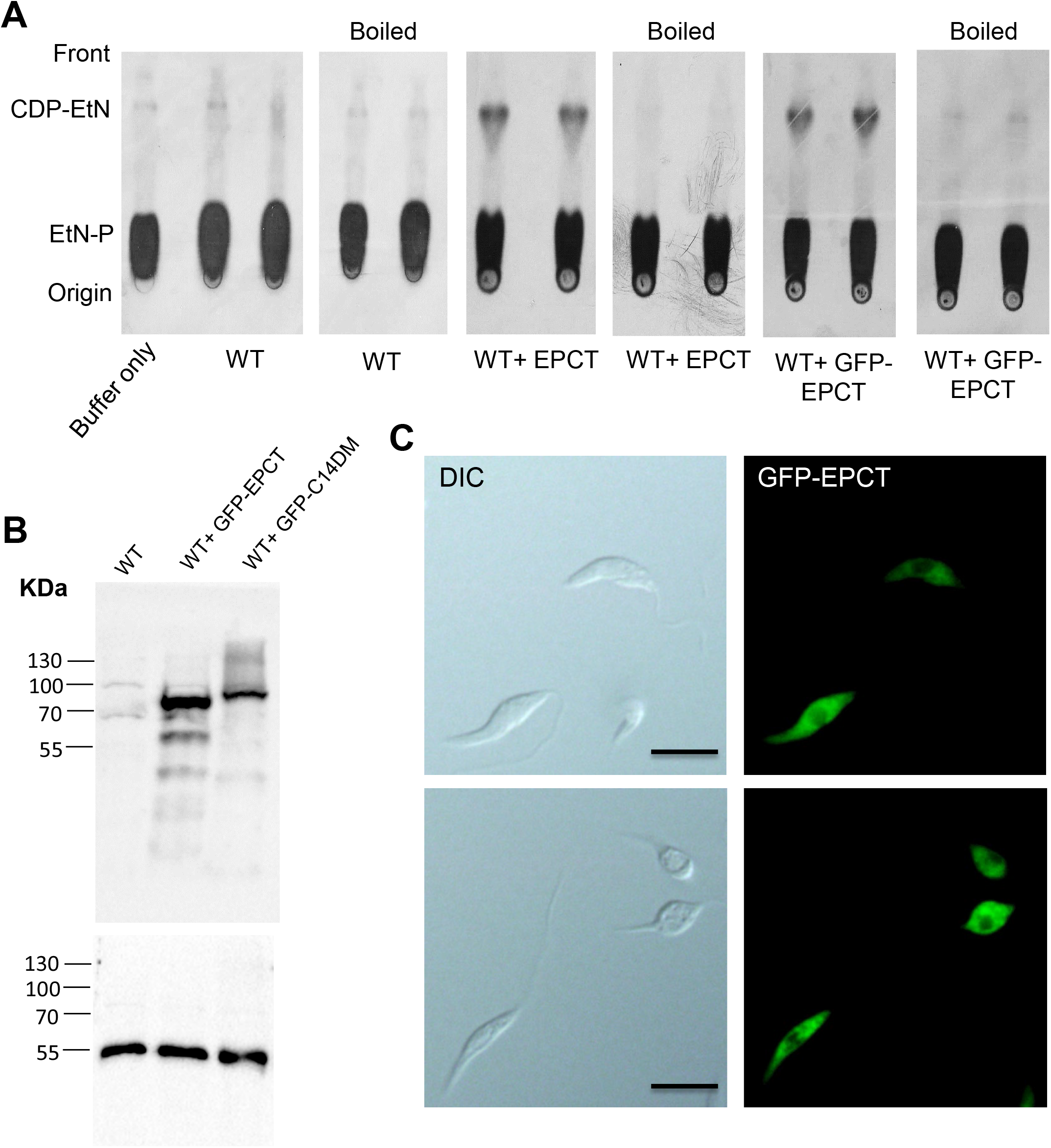
*L. major* EPCT is a functional enzyme located in the cytoplasm. (**A**) Whole cell lysates (boiled and unboiled, two repeats each) of WT, WT+EPCT, and WT+GFP-EPCT promastigotes were incubated with [^14^C]-EtN-P followed by TLC analysis as described in *Materials and Methods*. (**B**) Promastigote lysates of WT, WT+GFP-EPCT, and WT+C14DM-GFP (a control) were probed with an anti-GFP antibody (top) or anti-α-tubulin antibody (bottom). (**C**) Log phase (top) and stationary phase(bottom) promastigotes of WT+GFP-EPCT were examined by fluorescence microscopy. DIC: differential interference contrast. Scale bars: 10 µm.

### Chromosomal *EPCT*-null mutants cannot be generated without a complementing episome

To determine whether EPCT is required for survival in *L. major*, we first attempted to delete the two chromosomal *EPCT* alleles using the homologous recombination approach [41]. As demonstrated by Southern blot (Fig. S3A-B), we successfully replaced one *EPCT* allele with the blasticidin resistance gene (*BSD*) using this approach and generated several heterozygous *EPCT*+/- clones. However, repeated attempts to delete the second *EPCT* allele failed to recover any true knockouts (results from one attempt was shown in Fig. S3C-D), suggesting that a different method is needed to generate chromosomal *EPCT*-null mutants.

We then adopted the complementing episome-assisted knockout approach that has been used to study essential genes in *Leishmania* [42-44]. To do so, a pXNG4-EPCT plasmid (complementing episome) containing genes for *EPCT*, green fluorescence protein (*GFP*), nourseothricin resistance (*SAT*) and thymidine kinase (*TK*) was constructed and introduced into *EPCT*+/- parasites (*EPCT*+/- +pXNG4-EPCT), followed by attempts to delete the second chromosomal *EPCT* allele with the puromycin resistance gene (*PAC*) (Fig. S4). With this approach, we were able to generate multiple clones showing successful replacement of chromosomal *EPCT* with *BSD* and *PAC* (*epct*^−^+ pXNG4-EPCT in Fig. 3A-B and Fig. S4).

**Figure 3.**
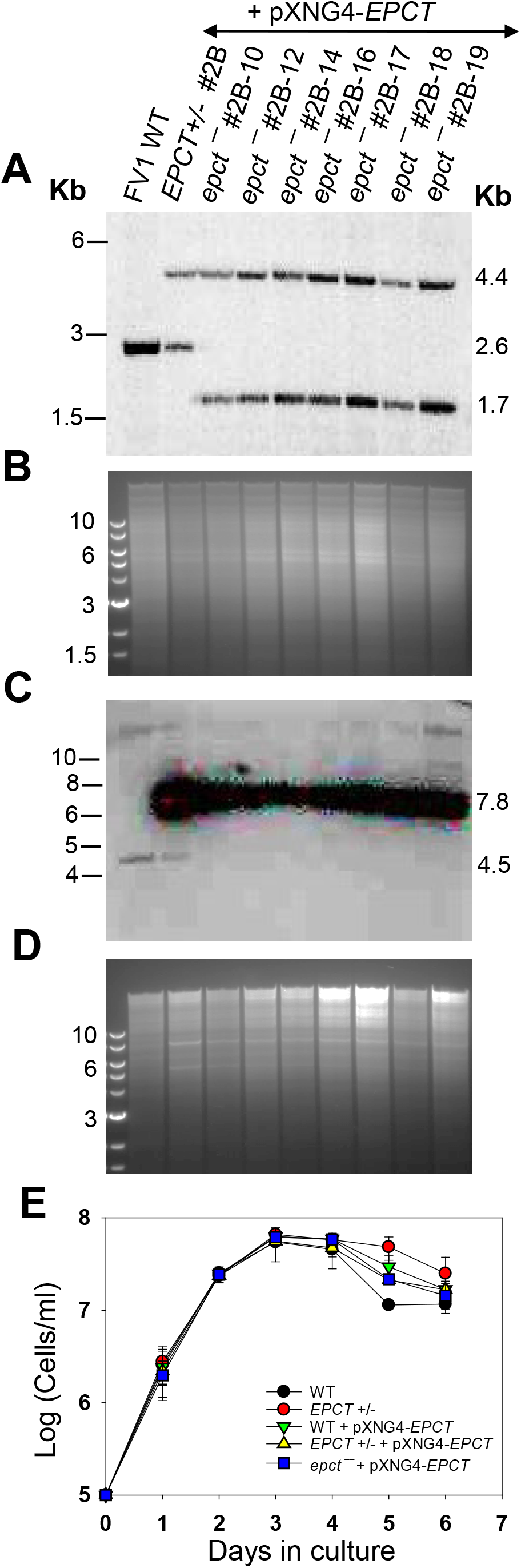
Generation of chromosomal *EPCT*-null mutants using an episome-assisted approach. (**A**-**D**) Genomic DNA samples from FV1 WT, *EPCT*+/- + pXNG4-*EPCT* (clone #2B), and *epct*^−^ + pXNG4-*EPCT* (seven clones) parasites were digested with *Aat* II (**A, B**) or *Kpn* I+*Avr* II (**C, D**) and analyzed by Southern blot using radiolabeled probes for an upstream flanking sequence (**A, B**) or the open reading frame of *EPCT* (**C, D**). DNA loading controls with ethidium bromide staining for **A** and **C** were included in **B** and **D**, respectively. The scheme of Southern blot and expected DNA fragment sizes were shown in Fig. S3. (**E**) Promastigotes of WT, *EPCT*+/- (#2), WT + pXNG4-*EPCT, EPCT+/−* + pXNG4-*EPCT* (clone #2B), and *epct*^−^ +pXNG4-*EPCT* (clone #2B-10) were cultivated at 27 °C in complete M199 media and culture densities were determined daily using a hemocytometer. Error bars indicate standard deviations from three biological repeats.

Southern blot with an *EPCT* ORF probe revealed high levels of pXNG4-EPCT plasmid (20-35 copies/cell) in these *epct*^−^+ pXNG4-EPCT parasites but no endogenous *EPCT* was detected (Fig. 3C-D and Fig. S4). Overexpression of *EPCT* from the pXNG4-EPCT plasmid did not have any significant effect on promastigote replication under normal culture conditions (Fig. 3E).

### EPCT is essential for *L. major* promastigotes

To evaluate the essentiality of EPCT, *EPCT*+/- +pXNG4-EPCT and *epct*^−^+ pXNG4-EPCT promastigotes were cultivated in the presence or absence of ganciclovir (GCV). Because of the *TK* expression from pXNG4-EPCT, adding GCV would trigger premature termination of DNA synthesis [42]. Thus, in the presence of GCV, parasites would favor the elimination of pXNG4-EPCT (which can be tracked by monitoring GFP fluorescence level) during replication to avoid toxicity, if the episome is dispensable [42]. As shown in Fig. 4A, after being cultivated in the presence of GCV for 14 consecutive passages, those chromosomal *EPCT*-null promastigotes still contained 60-74% of GFP-high cells, indicative of high pXNG4-EPCT retention levels after prolonged negative selection. In contrast, the pXNG4-EPCT plasmid was easily expelled from the *EPCT*+/-+pXNG4-EPCT parasites after 8 passages in GCV (Fig. 4A), suggesting that the remaining chromosomal *EPCT* allele made the episome expendable. Without GCV, we detected a gradual loss of episome in the *EPCT*+/-+pXNG4-EPCT but not *epct*^−^+ pXNG4-*EPCT* parasites (Fig. 4A), again arguing that the episome is indispensable in those chromosomal-null mutants. As controls, parasites grown in the presence of nourseothricin (the positive selection) consistently retained 85-91% GFP-high cells (“+SAT” in Fig. 4A-C).

**Figure 4.**
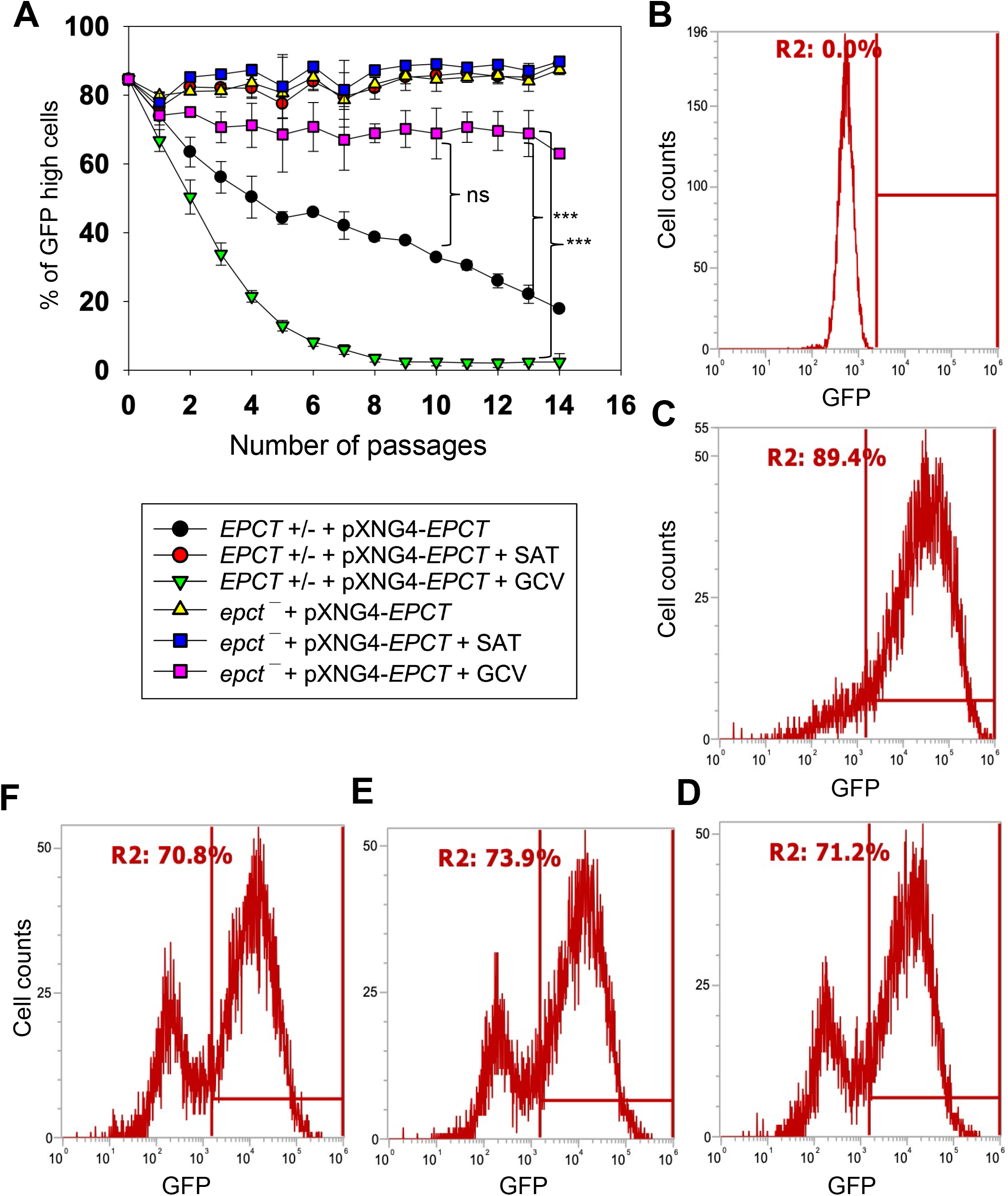
EPCT is indispensable during the promastigote stage. (**A**) Promastigotes were continuously cultivated in the presence or absence of ganciclovir (GCV) or nourseothricin (SAT) and passed every three days. Percentages of GFP-high cells were determined for every passage. Error bars represent standard deviations from three repeats (**: *p* < 0.01, ***: *p* < 0.001). (**B**-**D**) After 14 passages, WT (**B**) and *epct*^−^+pXNG4-EPCT parasites grown in the presence of SAT (**C**) or GCV (**D**) were analyzed by flow cytometry to determine the percentages of GFP-high cells (indicated by R2 in the histogram). (**E**-**F**) Two clones were isolated from the GFP-low population in **D** by sorting, amplified in the absence of SAT, and analyzed for GFP expression levels by flow cytometry.

Our analysis showed that 25-38% of *epct*^−^+ pXNG4-EPCT parasites became GFP-low after extended exposure to GCV (Fig. 4A and D). To examine whether these GFP-low cells were episome-free or harbored altered episome with mutations in *TK* and/or *GFP*, they were separated from GFP-high cells by fluorescence-activated cell sorting and individual clones were isolated after serial dilution. Two such clones were analyzed by flow cytometry showing similar percentages of GFP-high cells (70-74%) as the population prior to sorting (Fig. 4D-F), suggesting that the GFP-low cells were only viable in the presence of GFP-high cells.

Additionally, quantitative PCR (qPCR) analyses were performed on these clones using primers targeting the *GFP* region (to determine total plasmid copy number) and the *L. major* 28S rDNA (to determine total parasite number). Results revealed ∼8 copies of pXNG4-EPCT plasmid per cell in the *epct*^−^+ pXNG4-EPCT GCV-treated clones, whereas the *EPCT*+/- +pXNG4-EPCT clones isolated by the same process (GCV treatment → GFP-low sorting → serial dilution) contained <0.01 copy per cell (Fig. S5A). We also sequenced the pXNG4-EPCT plasmid DNA from GCV-treated *epct*^−^+ pXNG4-EPCT clones and did not find mutations in *TK, GFP* or *EPCT*. Finally, we found a significant growth delay in GCV-treated *epct*^−^+ pXNG4-EPCT but not *EPCT*+/-+pXNG4-EPCT promastigotes in comparison to control cells (Fig. S5B). This is consistent with the episome retention in *epct*^−^+ pXNG4-EPCT which leads to GCV-induced cytotoxicity. Together, these findings demonstrate the essential nature of EPCT in *L. major* promastigotes.

### EPCT is indispensable for *L. major* amastigotes

To investigate if EPCT is required for the survival of intracellular amastigotes, we infected BALB/c mice in the footpad with stationary phase promastigotes. After infection, half of the mice received daily GCV treatment, and the other half received equivalent amount of PBS (control group) for 14 consecutive days as previously described [44]. No significant changes in mouse body weight or movement were detected from GCV treatment (Fig. S6A). Mice infected by WT and *EPCT*+/- parasites exhibited equally rapid development of footpad lesions that correlated with robust parasite replication, indicating that one chromosomal copy of *EPCT* is sufficient for the mammalian stage (Fig. 5A-B). Importantly, mice infected by *EPCT*+/- +pXNG4-EPCT or *epct*^−^+ pXNG4-EPCT showed a 9-12-week delay in lesion development (Fig. 5A), and the delay was consistent with the significantly reduced parasite growth as determined by limiting dilution assay and qPCR (Fig. 5B and Fig. S6B). While GCV treatment had little impact on infections caused by WT, *EPCT*+/-, or *EPCT*+/- +pXNG4-EPCT parasites, it caused an additional 3-4-week delay in lesion progression for *epct*^−^+ pXNG4-EPCT (Fig. 5A-B), suggesting that GCV-induced cytotoxicity is more significant in the chromosomal null mutants than others.

**Figure 5.**
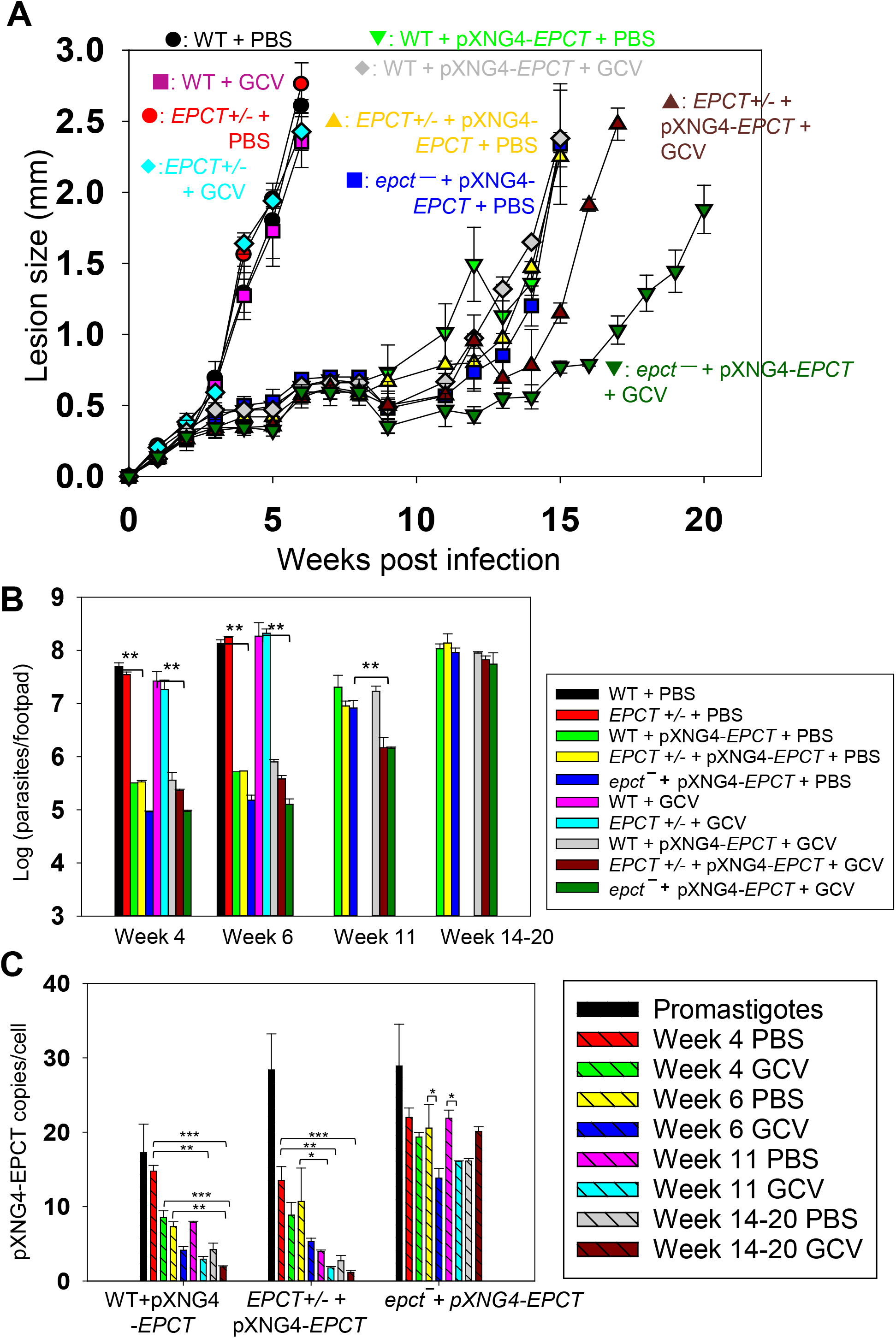
EPCT is indispensable during the amastigote stage. BALB/c mice were infected in the footpad with stationary phase promastigotes and treated with GCV or PBS as described in *Materials and Methods*. (**A**) Sizes of footpad lesions for infected mice were measured weekly. (**B**) Parasite numbers in infected footpads were determined by limiting dilution assay at the indicated times. Data for WT and *EPCT*+/- amastigotes were not available post 8 weeks when those infected mice had reached the humane endpoint. (**C**) The pXNG4-EPCT plasmid copy numbers in promastigotes and amastigotes (#/cell ± SDs) were determined by qPCR. Error bars represent standard deviations from three repeats (*: *p* < 0.05, **: *p* < 0.01, ***: *p* < 0.001).

To determine the pXNG4-EPCT copy number in amastigotes, we performed qPCR analyses on genomic DNA extracted from lesion-derived amastigotes. The average plasmid copy number per cell was revealed by dividing the total plasmid copy number with total parasite number. As shown in Fig. 5C, *EPCT*+/- +pXNG4-EPCT amastigotes contained 8-13 copies per cell at week 4 post infection and that number gradually went down to 1.5-2.5 copies per cell by week 14-20; and GCV treatment caused reduction in plasmid copy number as expected. The reduction in plasmid copy number over time suggests that the episome is not required in *EPCT*+/- +pXNG4-EPCT amastigotes. In comparison, *epct*^−^+ pXNG4-EPCT amastigotes maintained 15-30 plasmid copies per cell throughout the course of infection even with GCV treatment (Fig. 5C). Thus, these chromosomal *EPCT*-null amastigotes must retain a high episome level to survive which is consistent with their increased sensitivity to GCV (Fig. 5A-B), Together, these data argue that EPCT is critically important for the survival of intracellular amastigotes.

### EPCT overexpression leads to significantly attenuated virulence in mice

Results from our mouse experiments suggest that elevated EPCT expression from pXNG4-EPCT, a high copy number plasmid (Fig. 3C and Fig. 5C), may be responsible for the delayed lesion progression and amastigote growth from *EPCT*+/- +pXNG4-EPCT and *epct*^−^+ pXNG4-EPCT promastigotes (Fig. 5 and Fig. S6). To test this hypothesis, we introduced pXNG4-EPCT into WT parasites and the resulting WT +pXNG4-EPCT cells showed similar infectivity in mice as *EPCT*+/- +pXNG4-EPCT or *epct*^−^+ pXNG4-EPCT parasites (Fig. 5), indicating that this plasmid can attenuate virulence in the WT background as well. In addition, the *EPCT* gene was cloned into a pGEM vector along with its 5’- and 3’-flanking sequences (∼1 Kb each) and the resulting pGEM-EPCT was introduced into *EPCT*+/-parasites. Like cells with pXNG4-EPCT, these *EPCT*+/-+ pGEM-EPCT parasites displayed significantly reduced virulence and growth in BALB/c mice (Fig. S7), proving that these defects were caused by EPCT overexpression and not restricted to the pXNG4 plasmid.

Like *EPCT*+/- +pXNG4-EPCT, the episome copy number in WT +pXNG4-EPCT amastigotes decreased from 14-16 per cell at week 4 post infection to 2-4 per cell in week 14-20 (Fig. 5C). Curiously, neither *EPCT*+/- +pXNG4-EPCT amastigotes nor WT +pXNG4-EPCT amastigotes could completely lose the episome like *EPCT*+/- +pXNG4-EPCT promastigotes in culture after GCV treatment (Fig. 4A and Fig. 5A). Our previous studies on farnesyl pyrophosphate synthase (FPPS) and CEPT have demonstrated that heterozygous mutants of those genes lost most of their respective episome (<0.2 copy per cell for *FPPS+/−* +pXNG4-FPPS and *CEPT+/−* +pXNG4*-*CEPT) within 6 weeks post infection [18, 44]. It is possible that the slow replication rates of *EPCT*+/- +pXNG4-EPCT and WT +pXNG4-EPCT amastigotes prevented complete plasmid loss.

In addition to plasmid copy number, we also examined EPCT transcript levels by reverse transcription-qPCR using α-tubulin transcript as the internal standard (Fig. 6). For WT parasites, EPCT mRNA level went down ∼60% from promastigotes to amastigotes, consistent with the relatively slow replication rate for amastigotes (Fig. 6A). As expected, the presence of pXNG4-EPCT resulted in a 10-18-fold increase in EPCT mRNA levels (comparing to WT cells) during the promastigote stage (Fig. 6B). While the overexpression was less pronounced during the mammalian stage, we still observed a 2-8-fold increase in EPCT mRNA in *epct*^−^+ pXNG4-EPCT amastigotes (week 14-20 post infection) in comparison to WT amastigotes (Fig. 6C).

**Figure 6.**
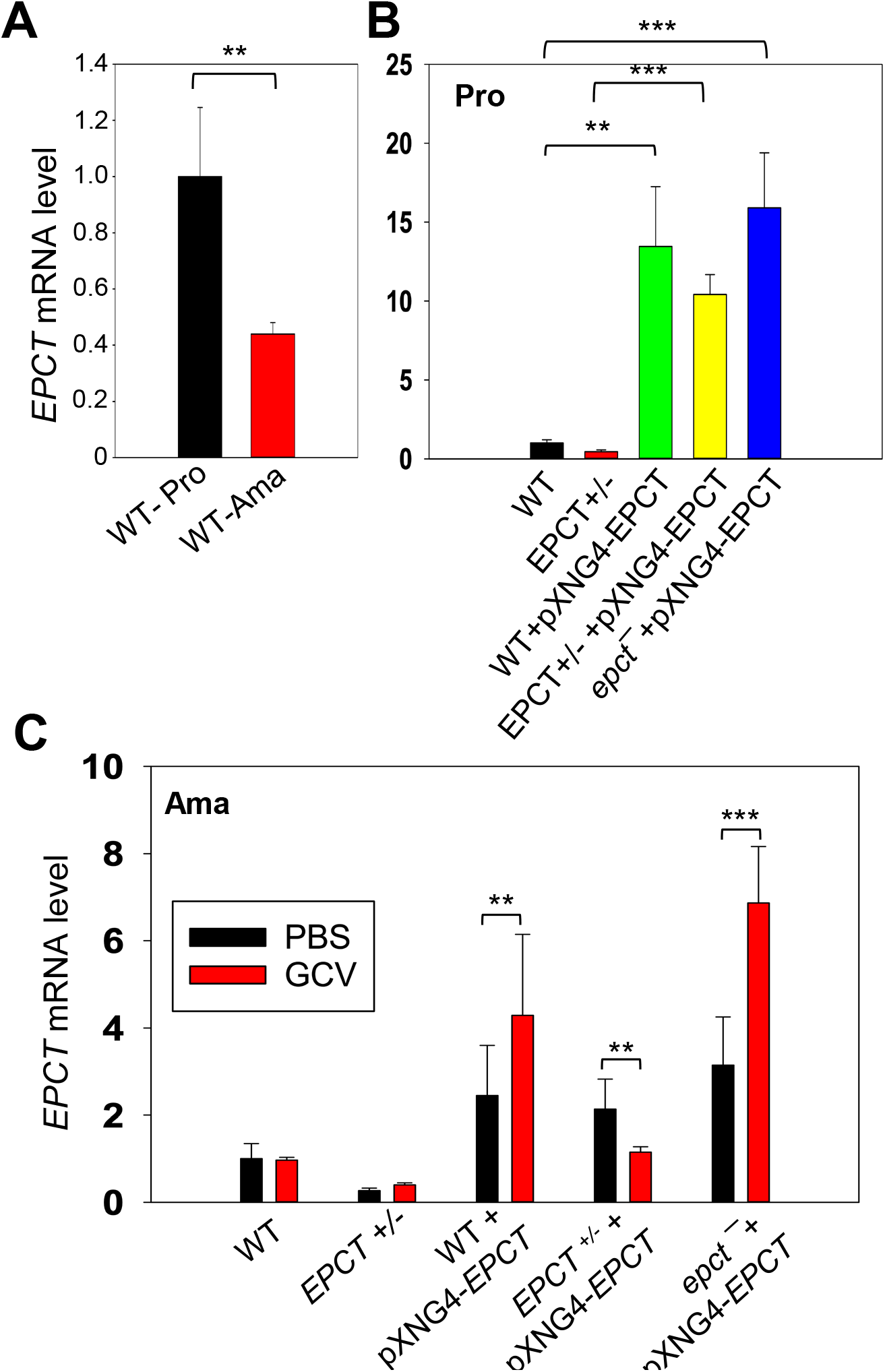
Expression level of *EPCT* mRNA in promastigotes and amastigotes. Total RNA was extracted from promastigotes (Pro) or amastigotes and the relative *EPCT* transcript levels were determined by qRT-PCR using the ΔΔCt method with α-tubulin gene being the internal control. (**A**) WT promastigotes and WT amastigotes (6 weeks post infection). (**B**) Promastigotes grown in the absence (WT and *EPCT*+/-) or presence of SAT (WT+pXNG4-EPCT, EPCT+/-+pXNG4-EPCT and *epct*^−^+pXNG4-EPCT). (**C**) Lesion derived amastigotes from mice treated with GCV or PBS (weeks 6 for WT and *EPCT*+/-, week 14-20 for EPCT overexpressors). Error bars represent standard deviation from three independent repeats (**: *p* < 0.01, ***: *p* < 0.001).

From these findings, we conclude that while EPCT is essential, episome-induced EPCT overexpression is detrimental to *L. major* growth during the mammalian stage.

### Changes in EPCT expression affects the syntheses of glycerophospholipids and GP63

To examine if *EPCT* expression level affects phospholipid composition in *L. major*, total lipids were extracted from stationary phase promastigotes and analyzed by electrospray ionization mass spectrometry (ESI-MS). To detect PME and diacyl-PE, we used precursor ion scan of m/z 196 in the negative ion mode and added 14:0/14:0-PE as a standard for quantitation (Fig. 7). As shown in Fig. 7, EPCT-overexpressing cells (WT +pXNG4-EPCT, EPCT+/-+pXNG4-EPCT and *epct*^−^+ pXNG4-EPCT) had 30-50% less PME (*p*18:0/18:2- and *p*18:0/18:1-PE) relative to WT promastigotes. Meanwhile, the overall PME level in EPCT+/-was only slightly less than WT (due to reduced level of *p*18:0/18:1-PE), and there was no significant change in the abundance of diacyl-PE molecules (Fig. 7). Similar ESI-MS analyses were performed to determine the levels of PC, phosphatidylinositol (PI), and IPC (the major sphingolipid in *Leishmania*). As summarized in Fig. 8 and Fig. S8, *EPCT* half knockout (EPCT+/-) or over-expression did not affect the composition of PC or IPC. Furthermore, *EPCT* overexpressing cells had 20-30% less 18:0/18:1-PI, the most abundant type of PI in comparison to WT cells (Fig. 9). We also detected a ∼70% reduction in the level of a18:0/18:1-PI (an alkyl-acyl PI) in WT+ pXNG4-*EPCT* parasites (Fig. 9). Collectively, these lipidomic analyses suggest that proper regulation of *EPCT* expression is vital for the balanced synthesis of glycerophospholipids.

**Figure 7.**
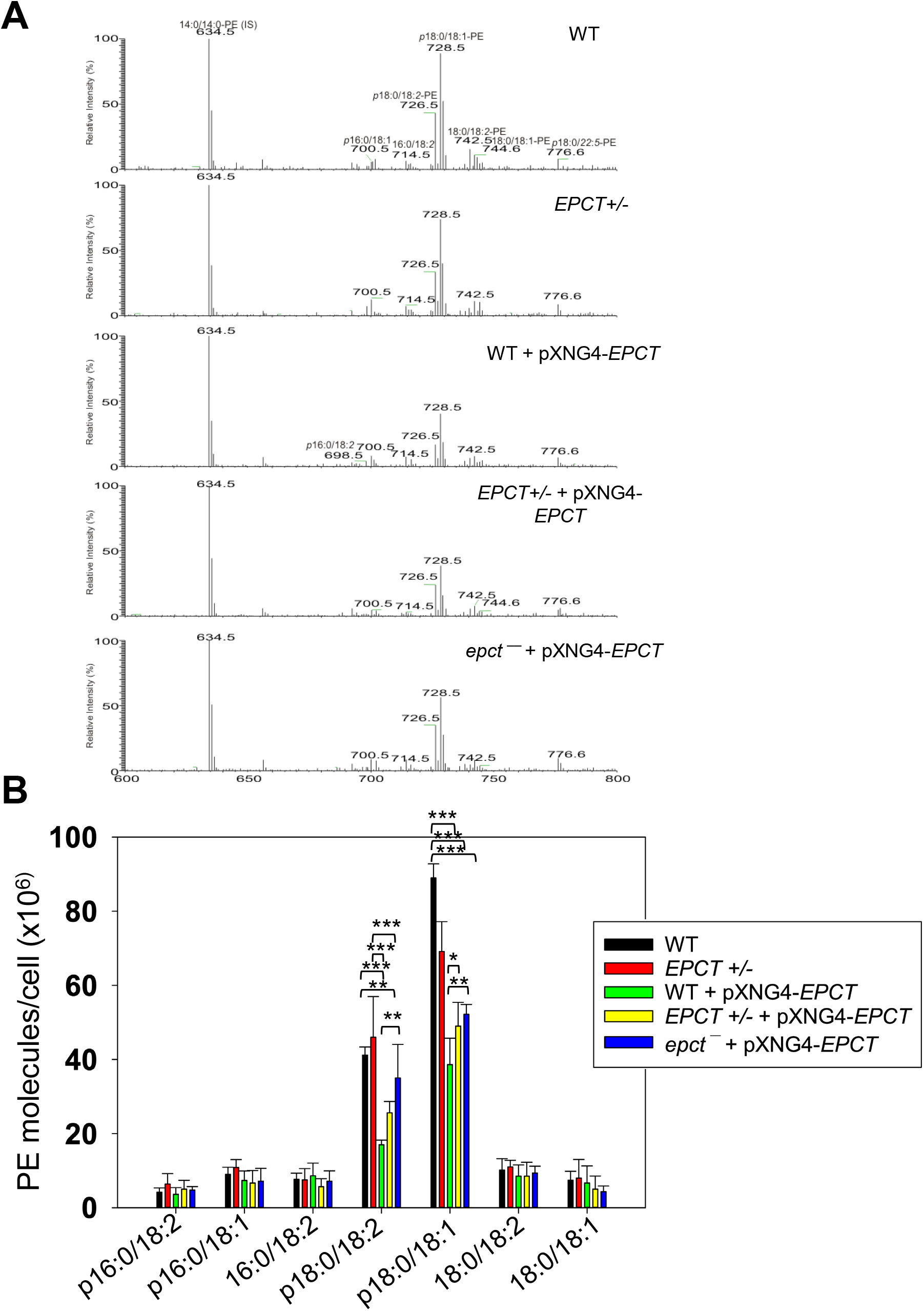
EPCT overexpression leads to reduced levels of PME. Total lipids were extracted from promastigotes and analyzed by ESI-MS in the negative ion mode (see text in the “Materials and Methods” for details). The 14:0/14:0-PE (m/z 634.5; [M – H]^-^ ion) was added as an internal standard. (**A**) Representative chromatograms of precursor ion scan of m/z 196 specifically monitoring PME and diacyl-PE species in a mixture. Common PME species such as *p*18:0/18:2-PE (m/z: 726.5) and *p*18:0/18:1-PE (m/z: 728.5) were indicated. (**B**) Plot of abundance of PME and diacyl PE lipids in different promastigote samples. Error bars represent standard deviation from 5 independent experiments (*: *p* < 0.05, **: *p* < 0.01, ***: *p* < 0.001).

**Figure 8.**
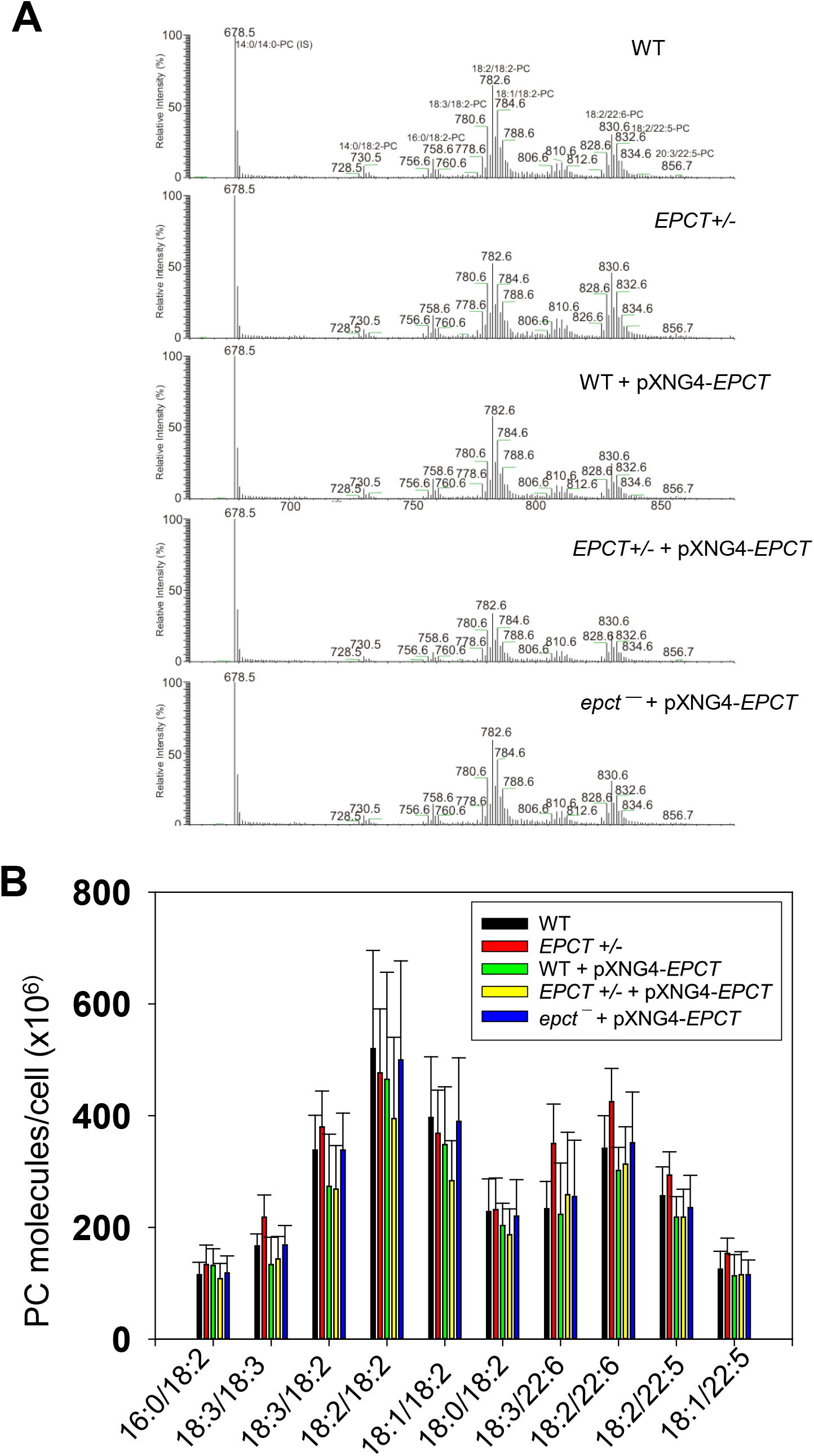
EPCT overexpression does not affect the synthesis of PC. Total lipids were extracted from promastigotes and analyzed by ESI-MS in the positive ion mode (see text in the “Materials and Methods” for details). The 14:0/14:0-PC (m/z 678.5; [M + H]^+^ ion) was added as an internal standard. (**A**) Representative chromatograms of precursor ion scan of m/z 184 specifically monitoring PC and sphingomyelin species in a mixture. The structures of common PC species were illustrated. (**B**) Plot of abundance of PC lipids in different promastigote samples. Error bars represent standard deviation from 5 independent experiments.

**Figure 9.**
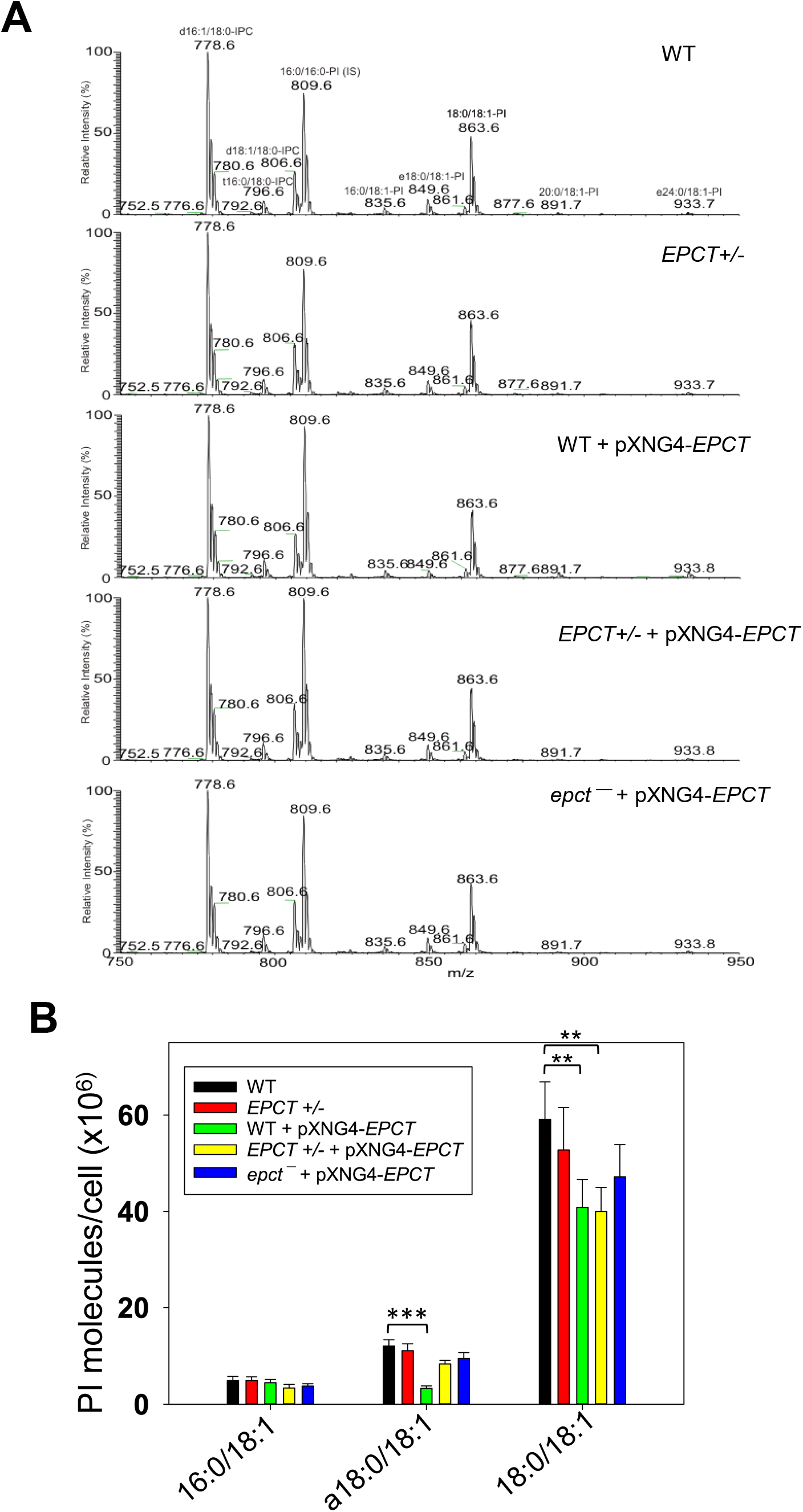
EPCT overexpression leads to reduced levels of PI. Total lipids were extracted from promastigotes and analyzed by ESI-MS in the negative ion mode. The 16:0/16:0-PI (m/z: 809.6) was added as an internal standard. (**A**) Representative chromatograms of precursor ion scan for m/z 241 (specific for PI). Common PI species were indicated. (**B**) Abundance of PI in promastigotes. Error bars represent standard deviation from 4 independent experiments (**: *p* < 0.01, ***: *p* < 0.001).

Next, because PE and PI are involved in the biosynthesis of GP63 (a zinc-dependent, GPI-anchored metalloprotease) and lipophosphoglycan (LPG), respectively, we examined whether *EPCT* under- or over-expression could influence the production of these surface glycoconjugates. Interestingly, immunofluorescence microscopy revealed a 4**-**5-fold reduction of GP63 in EPCT+/-, WT +pXNG4-EPCT, and EPCT+/-+pXNG4-EPCT parasites and a nearly 8-fold reduction in *epct*^−^ + pXNG4-EPCT (Fig. 10A and C). On the other hand, the cellular levels of LPG were unaltered in *EPCT* mutant lines (Fig. 10B and D). These findings argue that *ECPT* expression from its chromosomal locus are required for the proper synthesis of GP63.

**Figure 10.**
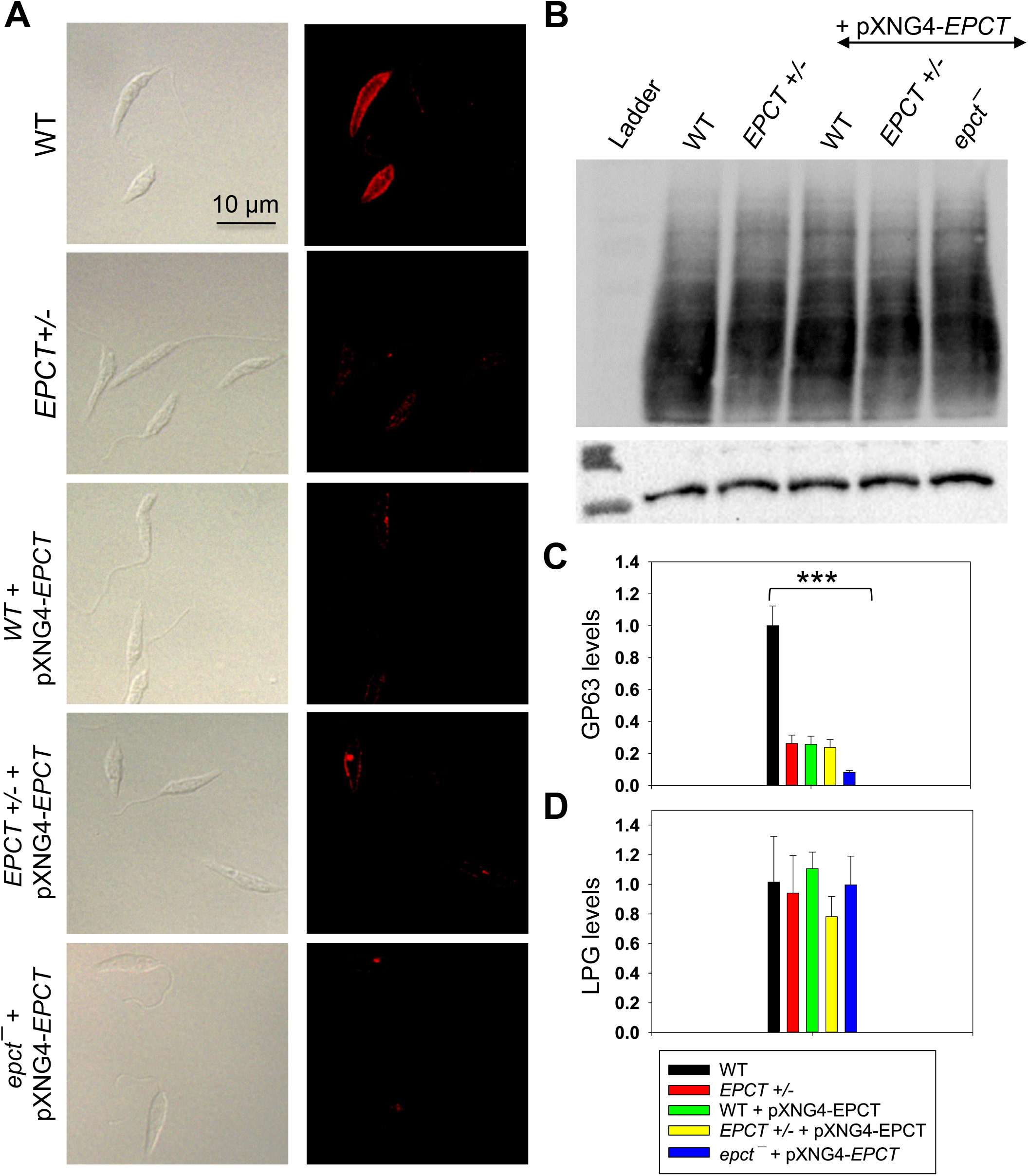
EPCT overexpression leads to reduced level of GP63 but not LPG. (**A**) WT, *EPCT*+/-, WT + pXNG4-*EPCT, EPCT+/-* + pXNG4-*EPCT*, and *epct***^−^** + pXNG4-*EPCT* promastigotes were labeled with an anti-GP63 monoclonal antibody followed by an anti-mouse IgG-Texas Red antibody (left: DIC, right: fluorescence). (**B**) Whole cell lysates from stationary phase promastigotes were analyzed by western blot using an anti-*L. major* LPG antibody (top) or anti-α-tubulin antibody (bottom). The relative abundances of GP63 (**C**) and LPG (**D**) were determined and normalized to WT levels. Error bars represent standard deviations from 3 independent repeats (***: *p* < 0.001).

### EPCT overexpression affects stress response

To further explore how EPCT overexpression may compromise *L. major* virulence, we examined the response of mutants to starvation, acidic pH and heat stress as tolerance to these stress conditions are essential for parasite survival in the macrophage phagolysosome. Under normal conditions (complete M199 medium, pH7.4, 27 °C), EPCT+/- and overexpressors proliferated at similar rates as WT promastigotes in log phase and exhibited good viability in stationary phase (Fig. 3E and Fig. 11A). However, if cells were transferred to phosphate-buffered saline (PBS, pH7.4) to test starvation tolerance, 50-60% of *EPCT* overexpressing cells (WT+pXNG4-EPCT, EPCT+/-+pXNG4-EPCT and *epct*^−^+ pXNG4-EPCT) died after 2-3 days in comparison to <20% death for WT parasites (Fig. 11B). These overexpressors also showed a 24–48-hour growth delay if they were cultivated in a pH5.0 medium (Fig. 11D). In addition, *epct*^−^+ pXNG4-*EPCT* parasites were slightly more sensitive to acidic stress (pH 5.0) and heat (37 °C) than WT in the stationary phase (Fig. 11C and F). We did not detect any significant difference in mitochondrial superoxide level between WT and EPCT mutants (Fig. S9). Whether these defects are linked to the altered lipid contents in EPCT overexpressors remains to be determined. Regardless, sensitivity to stress conditions likely contributes to their lack of virulence.

**Figure 11.**
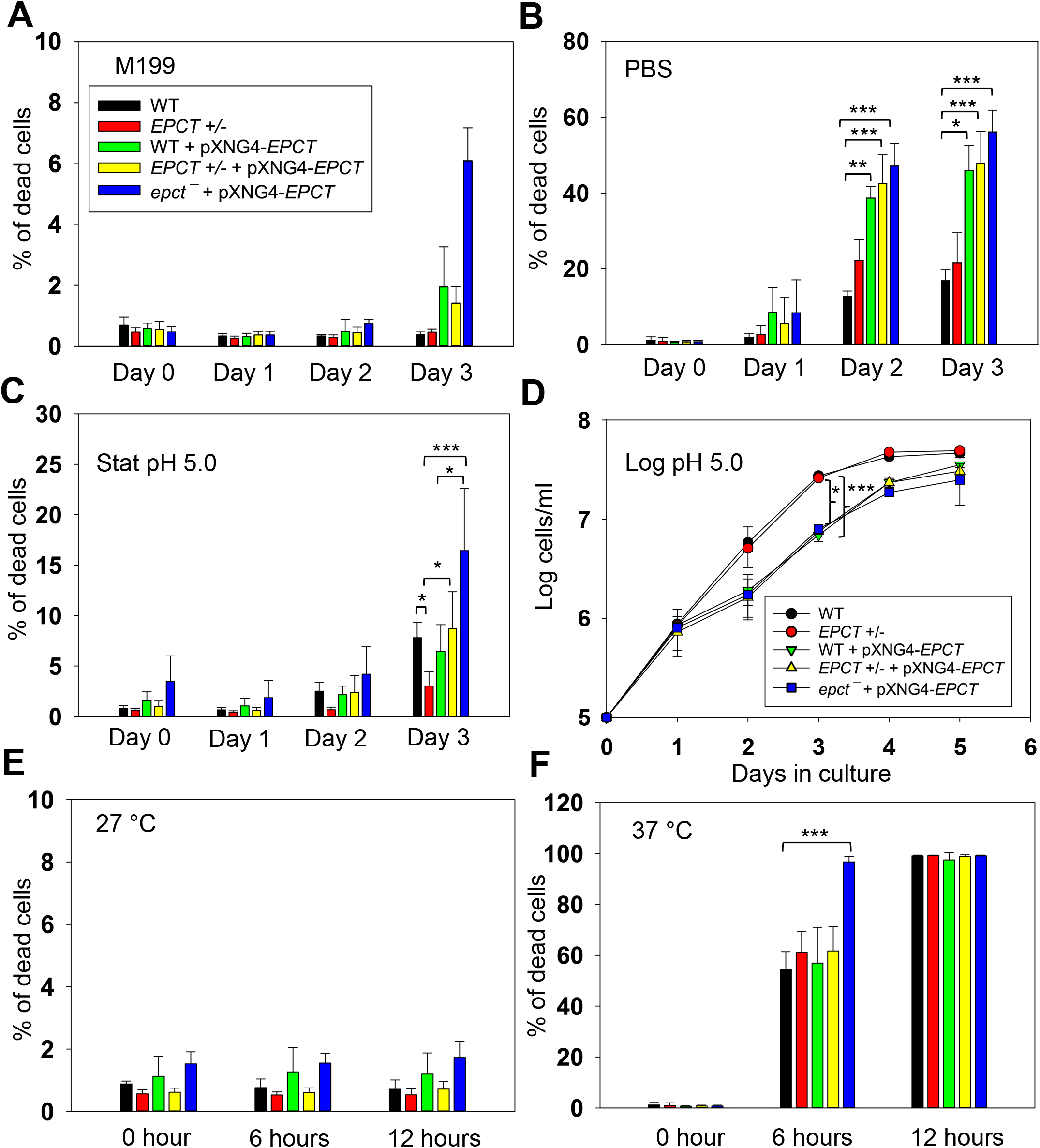
EPCT overexpression leads to heightened sensitivity to acidic pH and starvation conditions. (**A-B**) Promastigotes were cultivated in regular M199 media (pH 7.4) at 27 °C until reaching stationary phase. Cells were then kept in M199 media (**A**) or transferred into PBS (**B**). (**C-D**) Stationary phase (**C**) or log phase (**D**) promastigotes were cultivated in an acidic M199 medium (pH5.0). (**E-F**) Day 1 stationary phase promastigotes were inoculated in complete M199 media (pH 7.4) and incubated at 27 °C (**E**) or 37 °C (**F**). Percentage of dead cells were measured by flow cytometry after propidium iodide staining (**A**-**C, E** and **F**) and cell growth was determined by hemocytometer counting **(D**). Error bars represent standard deviations from 3-4 independent experiments (*: *p* < 0.05, **: *p* < 0.01, ***: *p* < 0.001).

## DISCUSSION

Using a complementing episome assisted knockout approach coupled with negative selection, we demonstrate that EPCT is indispensable throughout the life cycle in *L. major*. The same method was applied to verify the essentiality of several genes in *Leishmania* [42-44]. With EPCT being essential in both promastigotes and amastigotes, it is time to recognize EPCT as the most important enzyme of the Kennedy pathway. This is in sharp contrast to cholinephosphate cytidyltransferase (CPCT) which catalyzes the equivalent step in the choline branch of the Kennedy pathway by combing choline phosphate and CTP into CDP-choline (Fig. 1). *L*.

*major* CPCT-null mutants (*cpct*^−^) cannot incorporate choline into PC but contain similar levels of PC as WT parasites [45]. Loss of CPCT has no impact on promastigote growth in culture or their virulence in mice [45]. A similar study on CEPT which is directly responsible for the generation of diacyl-PE and PC revealed that this enzyme is only required for the promastigote but not amastigote stage [18]. These findings indicate that intracellular amastigotes can survive and proliferate without *de novo* PC synthesis by replying on the uptake/remodeling of host lipids.

Like with CEPT, chromosomal null mutants for *EPCT* cannot lose the complementing pXNG4-EPCT episome in culture, suggesting that *de novo* PE synthesis is required for the fast-replicating promastigotes (Fig. 4). The fact that EPCT is indispensable for amastigotes is intriguing, as amastigotes can acquire enough PC, which is far more abundant than PE [18]. One possibility is that the PE scavenging pathway, either direct uptake or remodeling of host lipids, is insufficient and amastigotes need certain level of *de novo* synthesis to meet the demand for PE. Alternatively, because of its central position in the Kennedy pathway, a loss of EPCT will not only abolish the *de novo* synthesis of PME and diacyl-PE, but also negatively affect the production of PC (from PE N-methylation) and PS (from PE interconversion). In comparison to EPT, CPCT and CEPT, disruption of EPCT would result in a more pleiotropic effect on the synthesis of multiple classes of glycerophospholipids (Fig. 1).

We observed a significant virulence attenuation from EPCT overexpression as parasites with pXNG4-ECPT or pGEM-EPCT showed delayed lesion progression and cell replication in mice (Fig. 5 and Fig. S7). Such defects were not found with the episomal overexpression EPT, CPCT, CEPT, or FPPS (an essential enzyme required for sterol synthesis) [14, 18, 44, 45]. As a cytosolic protein, EPCT generates CDP-EtN which is a substrate for EPT and CEPT to synthesize PME and diacyl-PE, respectively (Fig. 1 and 2). Both EPT and CEPT are localized in the endoplasmic reticulum (ER) [14, 18]. EPCT overexpression leads to reduced levels of PME but has little effect on diacyl-PE or PC (Fig. 7 and 8), suggesting that increased production of CDP-EtN is insufficient to boost PE synthesis and may instead cause substrate inhibition when substrate concentrations exceed the optimum level for certain enzymes which hinders product release [46]. It is not clear whether the altered PME synthesis in EPCT overexpressing cells is responsible for their heightened sensitivity to starvation or acidic pH. Our previous report on *L. major ept*^−^ mutants indicate that while these mutants are largely devoid of PME, they did not exhibit the same stress response defects as EPCT overexpressing cells [14]. It is possible that the increased cytosolic concentration of CDP-EtN, coupled with the depletion of EtN-P, causes cytotoxicity when parasite encounter starvation or acidic stress. We did not detect significant changes in mitochondrial superoxide production from EPCT under- or over-expression, suggesting that mitochondrial PE synthesis is not affected (Fig. S9).

While the alteration of *EPCT* expression had little impact on the cellular level of lipophosphoglycan (LPG), it did significantly reduce the expression of GP63, a zinc-dependent metalloprotease that plays pivotal role *Leishmania* infection through proteolytic cleavage of host complement proteins and subversion of macrophage signaling [47-50]. As illustrated in Fig. 10, *EPCT*+/- parasites contained 20-25% of WT-level GP63; episomal expression of *EPCT* did not rescue this defect and *epct*^−^+ pXNG4-EPCT cells had only 8-10% of WT-level GP63. These results suggest that *EPCT* must be expressed from its endogenous locus to fully support GP63 synthesis, during which PE is used as a donor to generate the EtN-P linkage between protein and GPI-anchor [7, 51].

In summary, our study establishes EPCT as the most important enzyme in the Kennedy pathway. It is crucial for *L. major* survival during the promastigote and amastigote stages.

Overexpression of EPCT alters lipid homeostasis and stress response, leading to severely attenuated virulence. Future studies will investigate how intracellular amastigotes balance *de novo* synthesis with scavenging to optimize their long-term survival and evaluate the potential of EPCT as a new anti-*Leishmania* target.

## MATERIALS AND METHODS

### Materials

Lipid standards for mass spectrometry studies including 1,2-dimyristoyl-sn-glycero-3-phosphoethanolamine (14:0/14:0-PE), 1,2-dimyristoyl-sn-glycero-3-phosphocholine (14:0/14:0-PC), and 1,2-dipalmitoyl-sn-glycero-3-phosphoinositol (16:0/16:0-PI) were purchased from Avanti Polar Lipids (Alabaster, AL). For the EPCT activity assay, phosphoryl ethanolamine [1,2-^14^C] (50-60 mCi/mmol) or [^14^-C] EtN-P was purchased from American Radiolabeled Chemicals (St Louis, MO). All other reagents were purchased from Thermo Fisher Scientific or Sigma Aldrich Inc unless otherwise specified.

### *Leishmania* culture and genetic manipulations

*Leishmania major* strain FV1 (MHOM/IL/81/Friedlin) promastigotes were cultivated at 27 °C in a complete M199 medium (M199 with 10% heat inactivated fetal bovine serum and other supplements, pH 7.4) [52]. To monitor growth, culture densities were measured daily by hemocytometer counting. Log phase promastigotes refer to replicative parasites at densities <1.0 × 10^7^ cells/ml, and stationary phase promastigotes refer to non-replicative parasites >2.0 × 10^7^ cells/ml.

To delete chromosomal *EPCT* alleles, the upstream and downstream flanking sequences (∼1 Kb each) of *EPCT* were amplified by PCR and cloned in the pUC18 vector. Genes conferring resistance to blasticidin (*BSD*) and puromycin (*PAC*) were then cloned between the upstream and downstream flanking sequences to generate pUC18-KO-EPCT:BSD and pUC18-KO-EPCT:PAC, respectively. To generate the *EPCT*+/- heterozygotes (*∆EPCT::BSD/EPCT*), wild type (WT) *L. major* promastigotes were transfected with linearized *BSD* knockout fragment (derived from pUC18-KO-EPCT:BSD) by electroporation and transfectants showing resistance to blasticidin were selected and later confirmed to be *EPCT*+/- by Southern blot as previously described [18]. To delete the second chromosomal allele of *EPCT*, we used an episome assisted approach as previously described [42]. First, the *EPCT* open reading frame (ORF) was cloned into the pXNG4 vector to generate pXNG4-EPCT and the resulting plasmid was introduced into *EPCT*+/- parasites. The resulting *EPCT*+/- +pXNG4-EPCT cell lines were then transfected with linearized *PAC* knockout fragment (derived from pUC18-KO-EPCT:BSD) and selected with 15 μg/ml of blasticidin, 15 μg/ml of puromycin and 150 μg/ml of nourseothricin. The resulting *EPCT* chromosomal null mutants with pXNG4-EPCT (*∆EPCT::BSD/∆EPCT::PAC* + pXNG4-EPCT or *epct*^−^+ pXNG4-EPCT) were validated by Southern blot as described in Fig. 2, Fig. S2 and S3. For EPCT activity assay and localization studies, the *EPCT* ORF was into the pXG vector or pXG-GFP’ vector to generate pXG-EPCT or pXG-GFP-EPCT respectively, followed by their introduction into WT *L. major* promastigotes by electroporation. Finally, a pGEM-5’-Phleo-DST IR-EPCT-3’ construct was generated by cloning the *EPCT* ORF along with its upstream- and downstream flanking sequences (∼1 Kb each) into a pGEM vector [53]. The resulting plasmid was introduced into *EPCT*+/- to drive *EPCT* overexpression in the presence of its flanking sequences.

### EPCT activity assay

EPCT assay was adopted based on a previously protocol [35]. Log phase promastigotes of WT, WT + pXG-EPCT, and WT + pXG-GFP-EPCT were resuspended in a lysis buffer based phosphate-buffered saline (PBS, pH 7.4) with 5 mM MgCl_2_, 0.1% Triton X-100, 5 mM DTT, and 1 X protease inhibitor cocktail at 4 × 10^8^ cells/ml. *Leishmania* lysate was incubated with 1 μCi of [^14^-C] EtN-P in a reaction buffer (100 mM Tris pH 7.5, 10 mM MgCl_2_, and 5 mM CTP) at room temperature for 20 min, boiled at 100 °C for 5 min to stop the reaction, and loaded directly on a Silica gel 60 TLC plate (10 μl each). TLC was developed in 100% ethanol:0.5% NaCl:25% ammonium hydroxide (10:10:1, v/v) and signals were detected via autoradiography at -80 °C. Mouse liver homogenate (with similar protein concentration as *Leishmania* lysate) was used as a positive control whereas boiled *Leishmania* lysate and mouse liver homogenate were included as negative controls. To determine the mobility of EPCT substrate (EtN-P) and product (CDP-EtN), non-radiolabeled EtN-P and CDP-EtN (40 nmol each) were loaded onto a Silica gel 60 TLC plate and processed with the same solvent as described above, followed by 0.2% ninhydrin spray.

### Fluorescence microscopy and western blot

An Olympus BX51 Upright Epifluorescence Microscope equipped with a digital camera was used to visualize the localization of GFP-EPCT and GP63 as previously described [14]. For GFP-EPCT, WT + pXG-GFP-EPCT promastigotes were attached to poly-L-lysine coated coverslips and fixed with 3.7% paraformaldehyde prior to visualization. GP63 staining of unpermeabilized parasites was performed using an anti-GP63 monoclonal antibody (#96/126, 1:500, Abcam Inc.) at ambient temperature for 30 min, followed by washing and incubation with a goat-anti-mouse-IgG-Texas Red antibody (1:500) for 30 min.

To determine the cellular level of lipophosphoglycan (LPG), promastigote lysates were boiled in SDS sample buffer at 95 °C for 5 min and resolved by SDS-PAGE. After transfer to PVDF membrane, blots were probed with mouse anti-LPG monoclonal antibody WIC79.3 (1:1000) [54] followed by goat anti-mouse IgG conjugated with HRP (1:2000). GFP-EPCT and GFP-C14DM were detected using a rabbit anti-GFP antibody (1:2000) followed by goat anti-rabbit IgG-HRP (1:2000). Antibody to alpha-tubulin was used as the loading control.

### Promastigote essentiality assay

WT, *EPCT*+/- +pXNG4-EPCT, and *epct*^−^+ pXNG4-EPCT promastigotes were inoculated in complete M199 media at 1.0 × 10^5^ cells/ml in the presence or absence of 50 μg/ml of GCV (the negative selection agent) or 150 μg/ml of nourseothricin (the positive selection agent). Every three days, cells were reinoculated into fresh media with the same negative or positive selection agents and percentages of GFP-high cells for each passage were determined by flow cytometry using an Attune NxT Acoustic Flow Cytometer. After 14 passages, individual clones of *EPCT*+/- +pXNG4-EPCT and *epct*^−^+ pXNG4-EPCT were isolated via serial dilution in 96-well plates. The GFP-high levels and pXNG4-EPCT plasmid copy numbers of selected clones were determined by flow cytometry and qPCR, respectively.

### Mouse footpad infection and *in vivo* GCV treatment

Female BALB/c mice (7-8 weeks old) were purchased from Charles River Laboratories International (Wilmington, MA). All animal procedures were performed as per approved protocol by Animal Care and Use Committee at Texas Tech University (PHS Approved Animal Welfare Assurance No A3629-01). To determine whether EPCT is required during the intracellular amastigote stage, day 3 stationary phase promastigotes were injected into the left hind footpad of BALB/c mice (1.0 × 10^6^ cells/mouse, 10 mice per group). For each group, starting from day one post infection, one half of the mice received GCV at 7.5 mg/kg/day for 14 consecutive days (0.5 ml each, intraperitoneal injection), while the other half (control group) received equivalent volume of sterile PBS. Footpad lesions were measured weekly using a Vernier caliper after anesthetization with isoflurane (air flow rate: 0.3-0.5 ml/hour). Mice were euthanized through controlled flow of CO_2_ asphyxiation when lesions reached 2.5 mm (humane endpoint) or upon the onset of secondary infections. Parasite numbers in infected footpads were determined by limiting dilution assay [55] and qPCR as described below.

### Quantitative PCR (qPCR) analysis

To determine parasite loads in infected footpads, genomic DNA was extracted from footpad homogenate and qPCR reactions were run in triplicates using primers targeting the 28S rRNA gene of *L. major* [18]. Cycle threshold (Ct) values were determined from melt curve analysis. A standard curve of Ct values was generated using serially diluted genomic DNA samples from *L. major* promastigotes (from 0.1 cell/reaction to 10^5^ cells/reaction) and Ct values >30 were considered negative. Parasite numbers in footpad samples were calculated from their Ct values using the standard curve. Control reactions included sterile water and DNA extracted from uninfected mouse liver.

To determine pXNG4-*EPCT* plasmid levels in promastigotes and lesion-derived amastigotes, a similar standard curve was generated using serially diluting pXNG4-EPCT plasmid DNA (from 0.1 copy/reaction to 10^5^ copies/reaction) and primers targeting the *GFP* region. qPCR was performed with the same set of primers on DNA samples and the average plasmid copy number per cell was determined by dividing total plasmid copy number with total parasite number based on Ct values.

To determine EPCT transcript levels, total RNA was extracted from promastigotes or lesion-derived amastigotes and converted into cDNA using a high-capacity reverse transcription kit (Bio-Rad), followed by qPCR using primers targeting *EPCT* or α-tubulin coding sequences. The relative expression level of *EPCT* was normalized to that of α-tubulin using the 2^−ΔΔ(Ct)^ method [56]. Control reactions were carried out without leishmanial RNA and without reverse transcriptase.

### Lipid extraction and mass spectrometric analyses

Total lipids from stationary phase promastigotes (1.0 × 10^8^ cells/sample) were extracted using the Bligh & Dyer method [57]. Commercial non-indigenous lipid standards were added to cell lysates as internal standards at the time of lipid extraction (1.0 × 10^8^ molecules/cell for 14:0/14:0-PE, 5.0 ×10^7^ molecules/cell for 14:0/14:0-PC, and 1.0 × 10^8^ molecules/cell for 16:0/16:0-PI). These lipid standards are absent from *L. major* promastigotes. Determination of lipid families by electrospray ionization mass spectrometry (ESI-MS) was carried out by a Thermo Vantage TSQ instrument applying precursor ion scan of m/z 196 for PE, precursor ion scan of m/z 241 for PI and IPC in the negative-ion mode, and precursor ion scan of m/z 184 for PC in the positive-ion mode [58]. Individual lipid species and their structures were also confirmed by high resolution mass spectrometry performed on a Thermo LTQ Obitrap Velos with a resolution of 100,000 (at m/z 400). All lipidomic analyses were performed five times.

### *Leishmania* stress response assays and determination of mitochondrial ROS level

For heat tolerance, *L. major* promastigotes grown in complete M199 media were incubated at either 27 °C or 37 °C. To test their sensitivity to acidic pH, promastigotes were transferred to a pH5.0 medium (same as the complete M199 medium except that the pH was adjusted to 5.0 using hydrochloric acid). For starvation response, promastigotes were transferred to PBS (pH 7.4). Cell viability was determined at the indicated times by flow cytometry after staining with 5 μg/ml of propidium iodide. Parasite growth was monitored by cell counting using a hemocytometer.

Mitochondrial superoxide accumulation was determined as described previously [34]. Briefly, log phase promastigotes were transferred to PBS (pH 7.4) and stained with 5 μM of MitoSox Red. After 25 min incubation at 27 °C, the mean fluorescence intensity (MFI) for each sample was measured by flow cytometry.

### Statistical analysis

Unless otherwise specified, experiments were repeated three to five times. Symbols or bars represent averaged values and error bars represent standard deviations. Differences between groups were assessed by one-way ANOVA (for three or more groups) using the Sigmaplot 13.0 software (Systat Software Inc, San Jose, CA) or student *t* test (for two groups). *P* values were grouped in all figures (***: *p* < 0.001, **: *p* < 0.01, *: *p* < 0.05).

## ACKNOWLEDGMENTS

This study was supported by NIH grants R15AI156746 (PI) and R01AI139198 (co-I) to KZ. Lipidomic analysis was supported by NIH grants P41-GM103422, P60-DK20579, P30-DK56341 to the Biomedical Mass Spectrometry Resource at Washington University in St. Louis.

## AUTHOR CONTRIBUTIONS

Conceptualization and Project Design: Somrita Basu, Kai Zhang

Data Generation: Somrita Basu, Mattie Pawlowic, Fong-Fu Hsu, Geoff Thomas

Formal data analysis: Somrita Basu, Kai Zhang, Fong-Fu Hsu

Investigation: Somrita Basu, Kai Zhang, Fong-Fu Hsu

Fund Acquisition: Kai Zhang, Fong-Fu Hsu

Writing: Somrita Basu, Kai Zhang

## SUPPORTING INFORMATION

**Figure S1.**
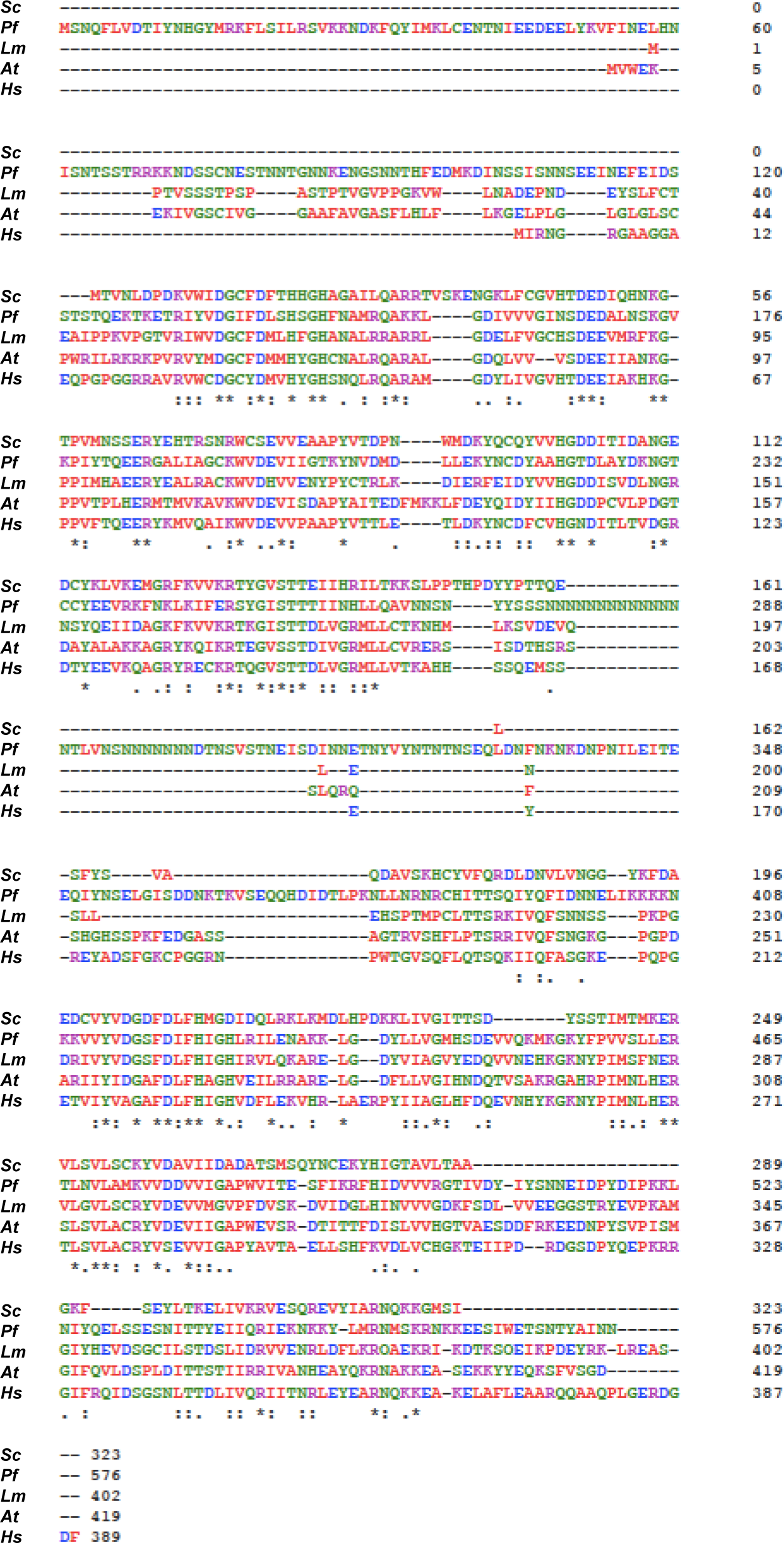
Alignment of EPCT amino acid sequences from *Saccharomyces cerevisiae* (*Sc*, Genbank: P33412), *Plasmodium falciparum* (*Pf*, Plasmodb: PfDd2_130053500), *Leishmania major* (*Lm*, Tritrypdb: LmjF.32.0890), *Arabidopsis thaliana* (*At*, TAIR: AT2G38670.1), and *Homo sapiens* (*Hs*, Genbank: Q99447) using Clustal Omega. Asterisks (*): fully conserved residues; colons (:) highly similar residues; periods (.): moderately similar residues. Color code for amino acids: red-nonpolar; green-polar; blue-acidic; purple-basic.

**Figure S2.**
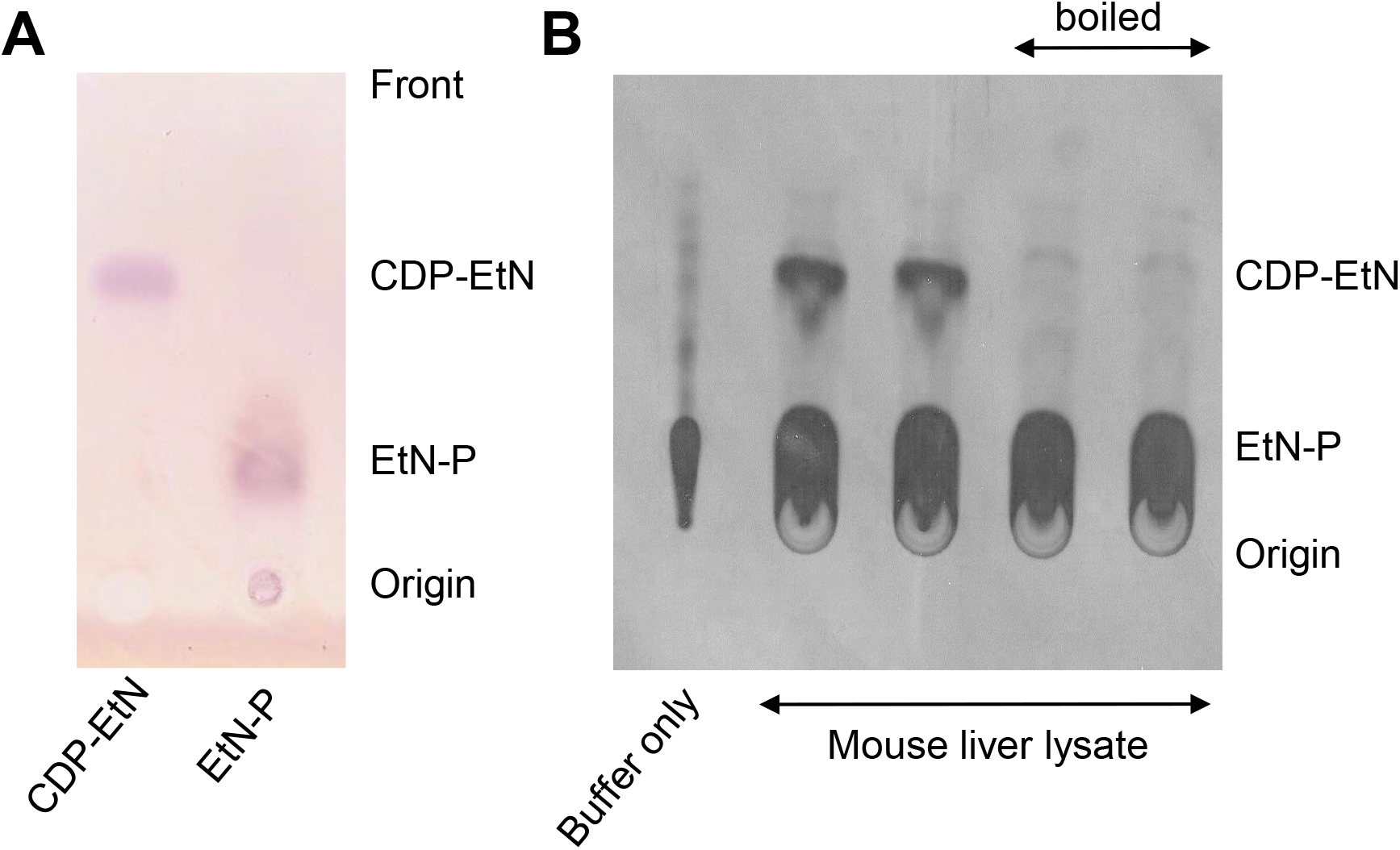
Detection of EPCT activity with thin layer chromatography (TLC). (**A**) EtN-P or CDP-EtN (40 nmol each) are resolved by TLC as described in *Materials and Methods*. The plate was dried and sprayed with 0.2% ninhydrin to show the positions of EtN-P and CDP-EtN. (**B**) Mouse liver lysates (boiled and un-boiled, two repeats each) were incubated with [^14^C]-EtN-P at room temperature, followed by TLC analysis and signals were detected by autoradiography.

**Figure S3.**
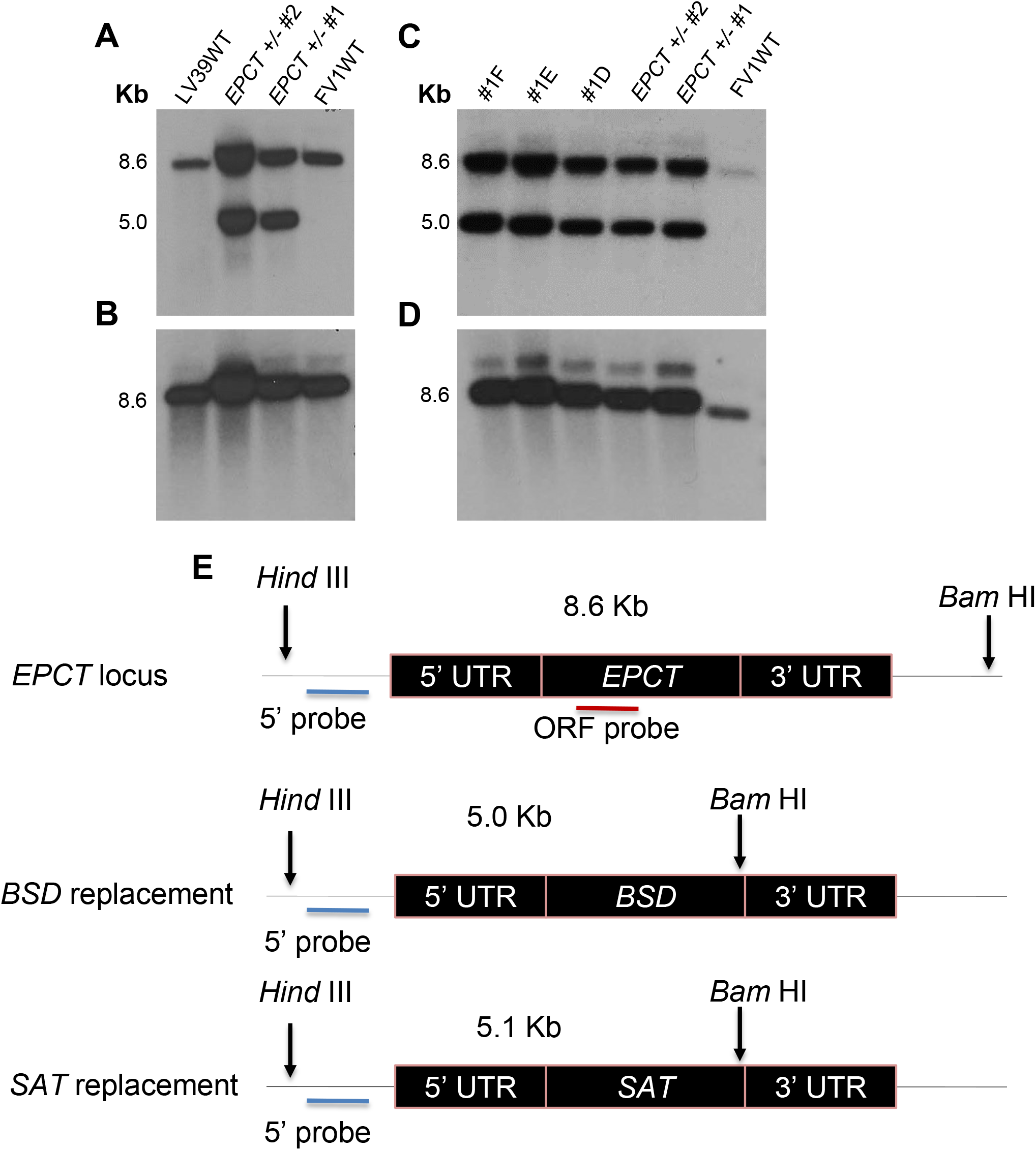
Chromosomal *EPCT-*null mutants cannot be generated without a complementing episome. Genomic DNA samples from *L. major* FV1 WT, LV39 WT, *EPCT*+/- (#1 and #2), and putative *epct*^−^ (#1D, #1E, and #1F) parasites were digested by *Hind* III + *Bam* HI followed by Southern blot analyses using radiolabeled probes for an upstream flanking sequence (5’ probe: **A** and **C**) and the open reading frame of *EPCT* (ORF probe: **B** and **D**). The approximate recognition sites of *Hind* III and *Bam* HI and expected DNA fragment sizes are indicated in **E**.

**Figure S4.**
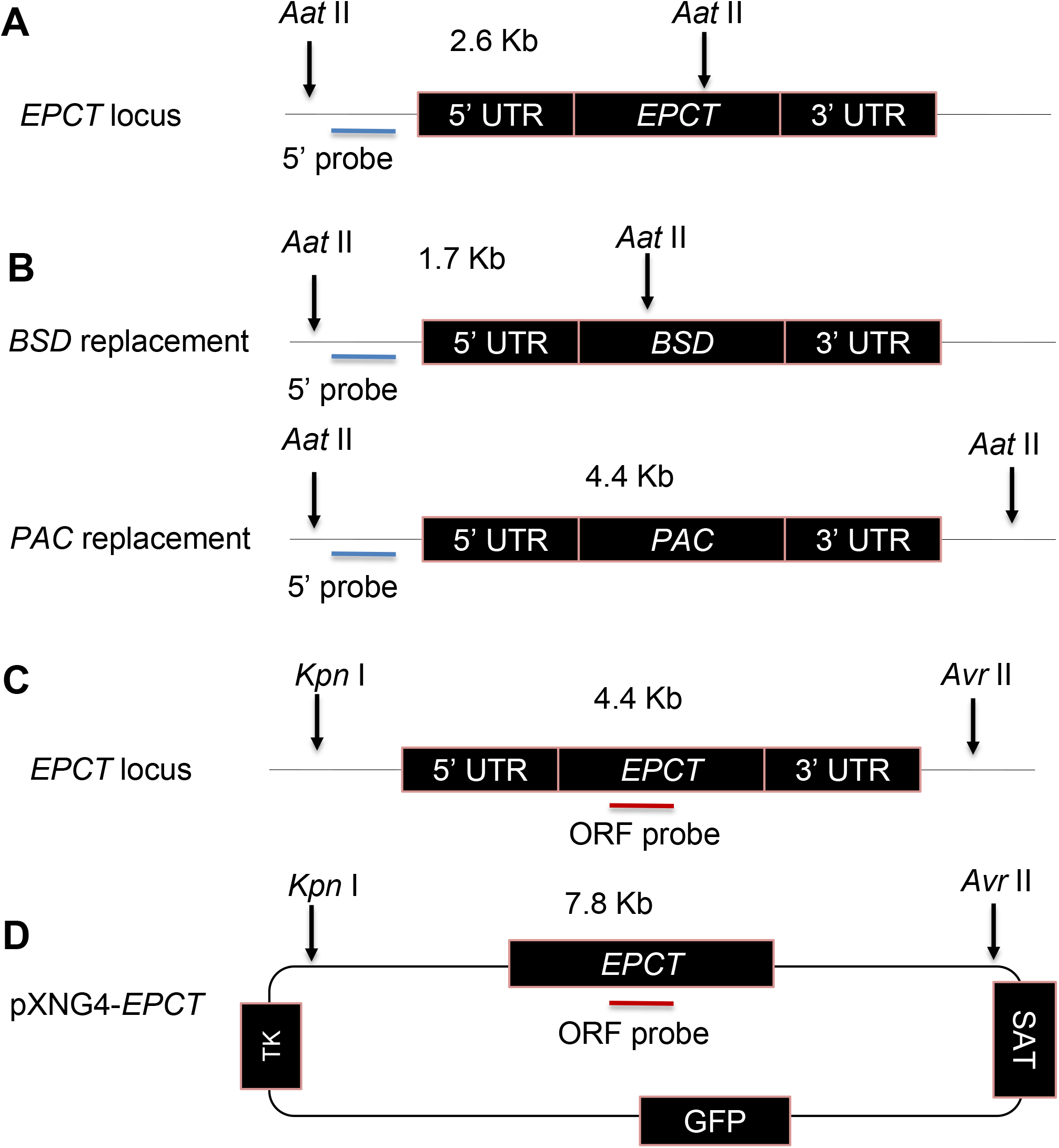
The scheme of Southern blot in Fig. 2. Expected DNA fragment sizes for using the 5’ probe (**A**-**B**) or ORF probe of *EPCT* (**C**-**D**) are indicated. TK: thymidine kinase, GFP: green fluorescent protein, SAT: nourseothricin resistance gene.

**Figure S5.**
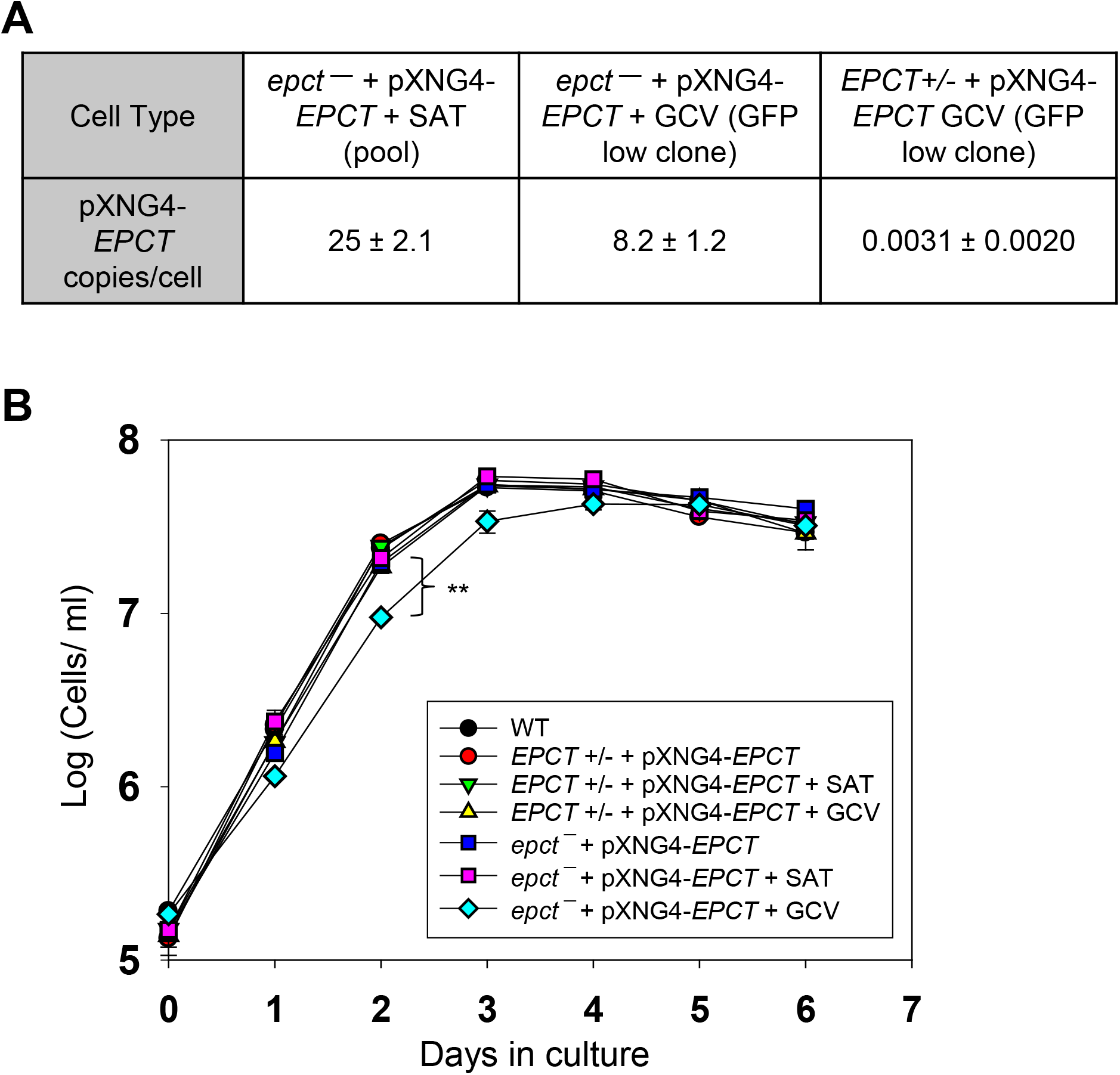
Chromosomal *EPCT*-null promastigotes cannot lose the pXNG4-EPCT episome. (**A**) *EPCT*+/- + pXNG4-EPCT and *epct*^−^ + pXNG4-EPCT promastigotes were cultivated in the presence of SAT or GCV for 14 passages (as pools) and individual clones were isolated via sorting followed by serial dilution. Plasmid copy number numbers (average ± SDs) were determined by qPCR. (**B**) Promastigotes were cultivated at 27 °C in complete M199 media and culture densities were determined daily using a hemocytometer. Error bars indicate standard deviations from three biological repeats (**: *p* < 0.01).

**Figure S6.**
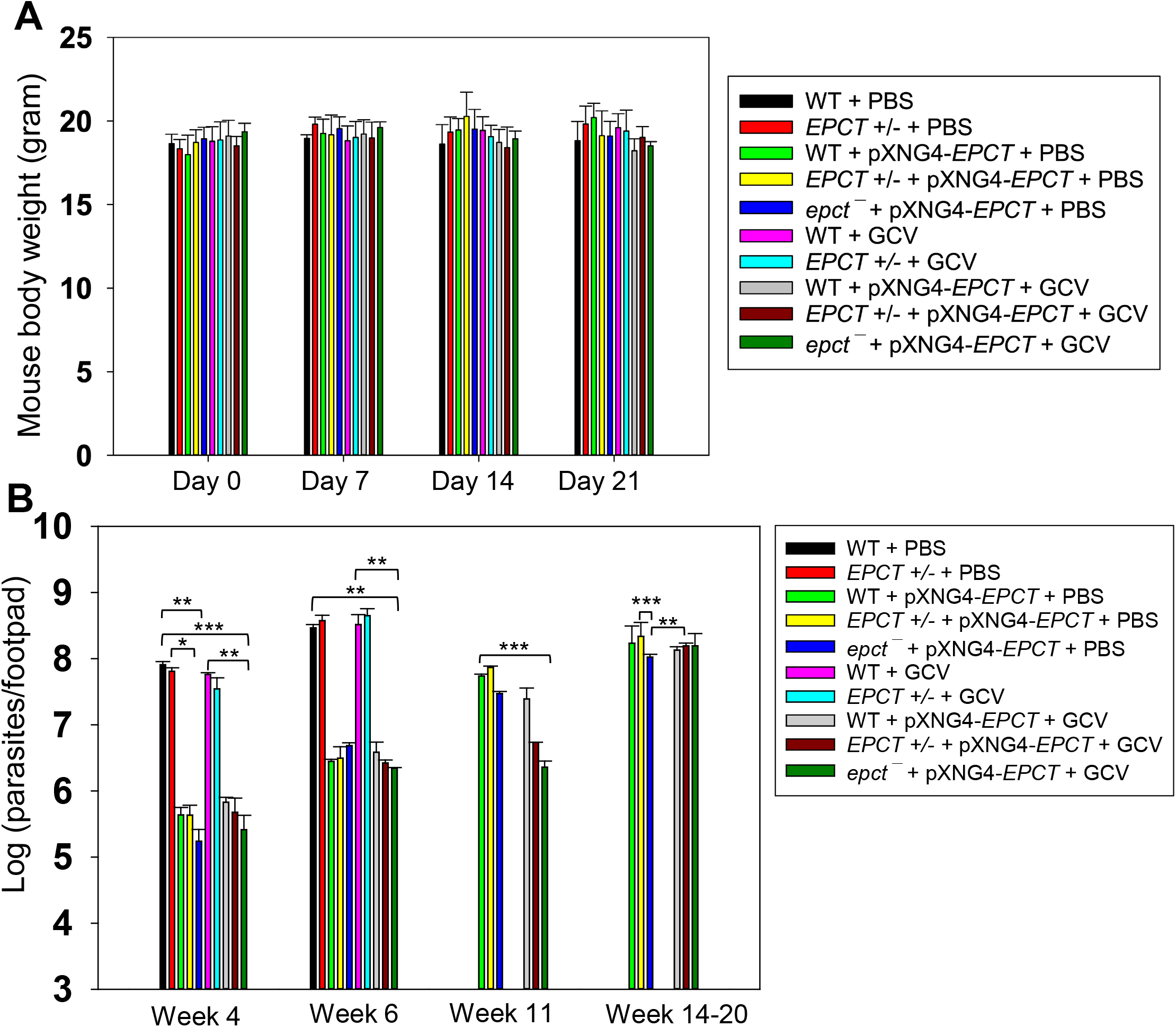
EPCT overexpression leads to reduced growth in BALB/c mice. Following footpad infection, mice were treated with GCV or PBS and euthanized at the indicated timepoints. (**A**) Mouse body weights were measured at day 0-21 post infection. (**B**) Genomic DNA samples were prepared from lesion-derived amastigotes and parasite loads were determined by qPCR using primers targeting the *L. major* 28S rDNA gene (*: *p* < 0.05, **: *p* < 0.01, ***: *p* < 0.001).

**Figure S7.**
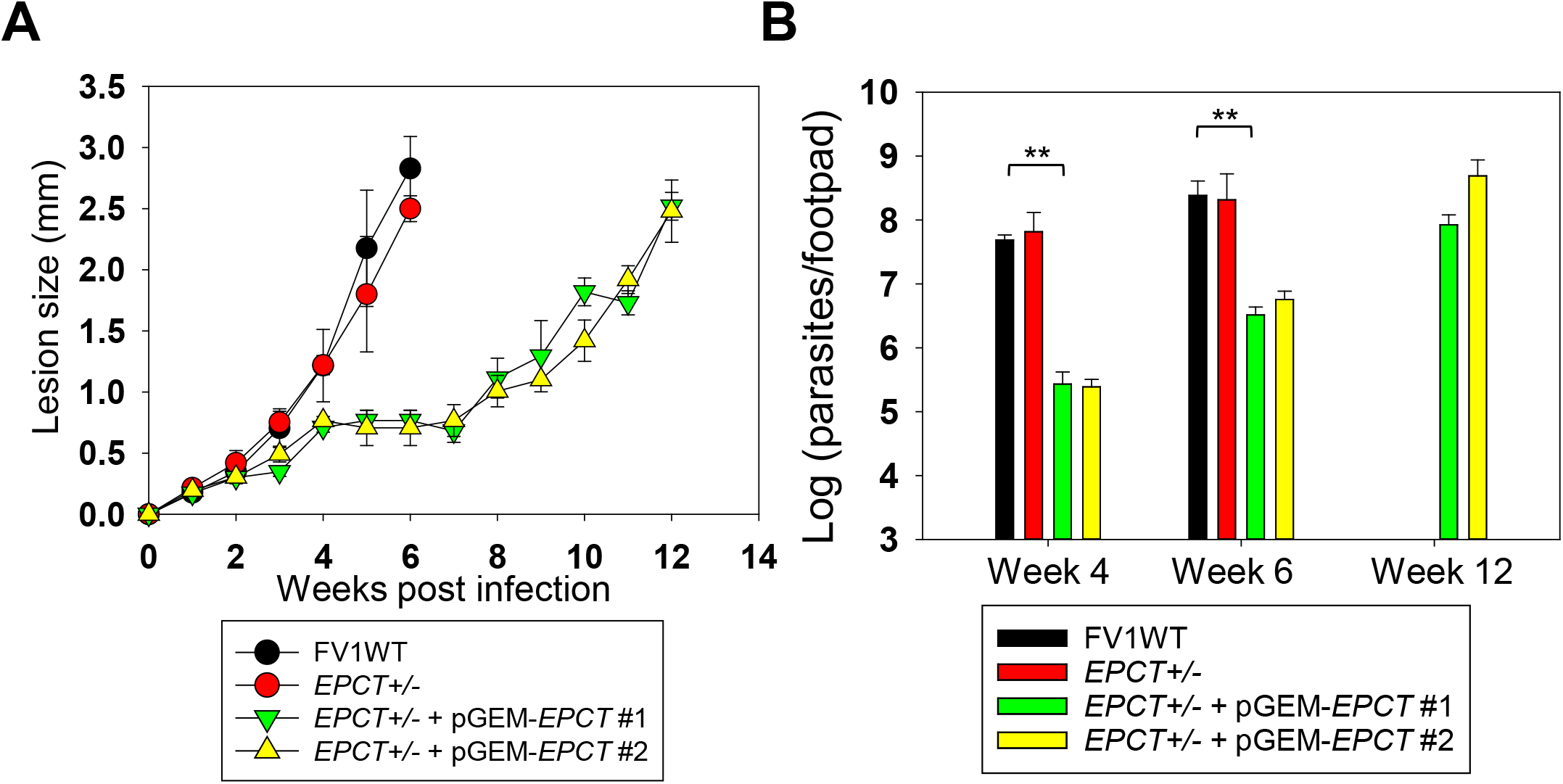
EPCT overexpression leads to attenuated virulence in BALB/c mice. Stationary phase promastigotes were injected into the footpad of BALB/c mice as described in *Materials and Methods*. Footpad lesion sizes were measured using a Vernier caliper (**A**) and parasite loads were determined by qPCR (**B**). **: *p* < 0.01.

**Figure S8.**
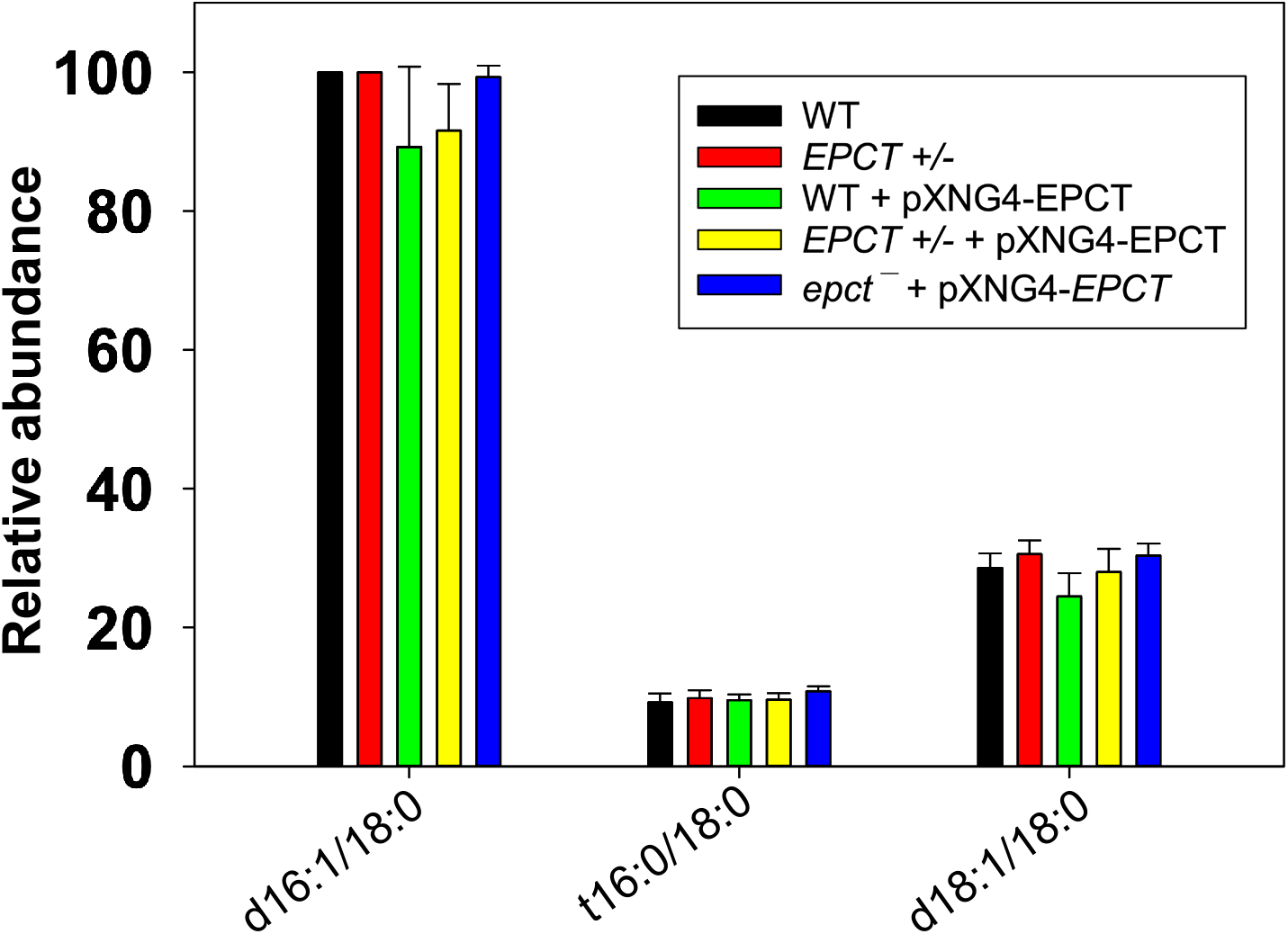
EPCT overexpression does not affect the cellular levels of IPC. Total lipids were extracted from stationary phase promastigotes and analyzed by ESI/MS in the negative ion mode using both total ion current scan and precursor ion scan of m/z 241. Error bars represent standard deviations from 4 independent experiments.

**Figure S9.**
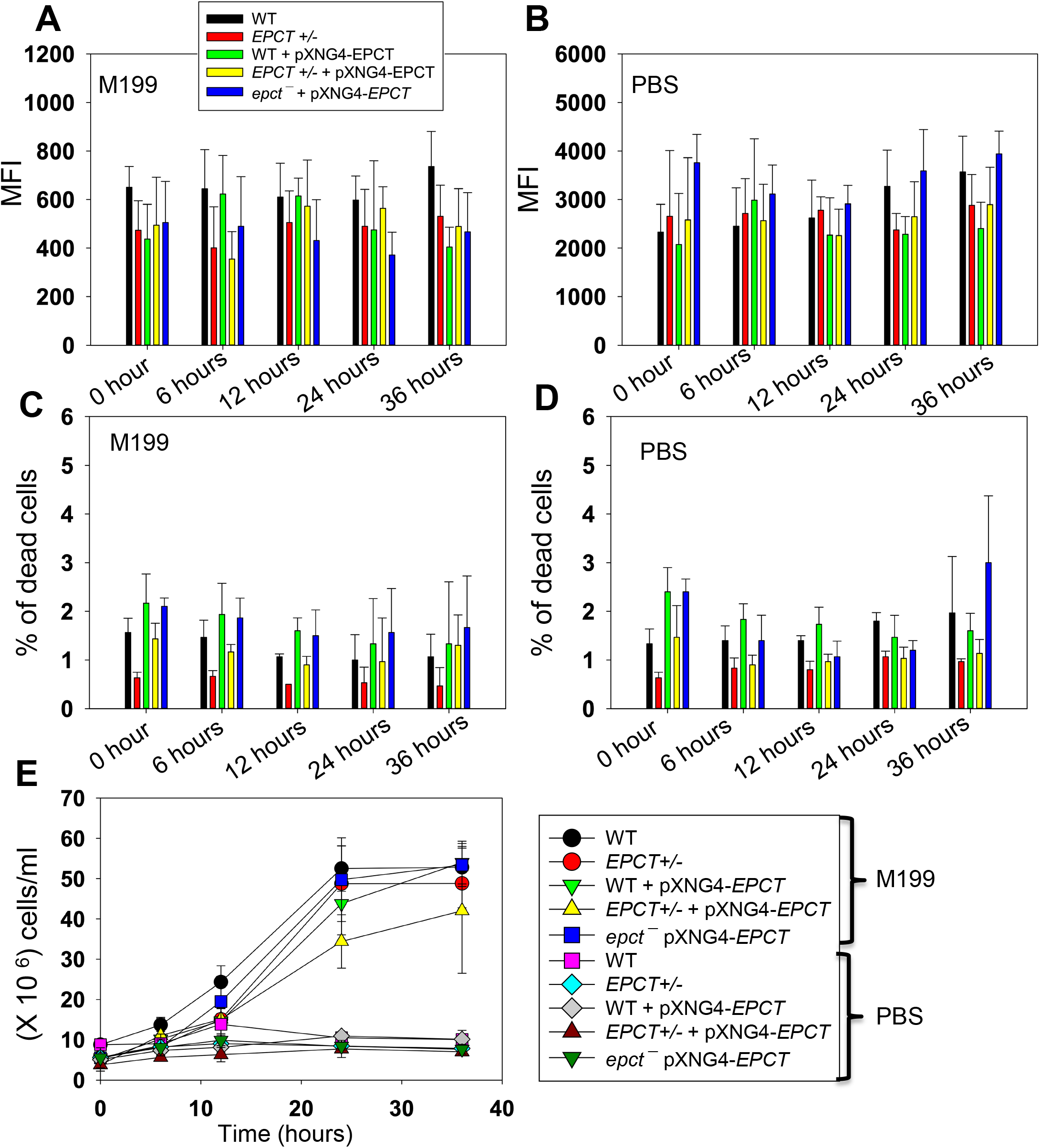
EPCT overexpression does not affect mitochondrial ROS production. Log phase promastigotes were cultivated in complete M199 medium **(A, C)** or transferred to PBS **(B, D)** and labeled with MitoSox Red for 25 min. Mean fluorescence intensity (MFI) for MitoSox Red (**A, B**) and percentages of dead cells (**C, D**) were determined by flow cytometry at the indicated timepoints. Cell growth rates were determined by hemocytometer counting **(E)**. Error bars represent standard deviations from three independent experiments.

**Table S1.**
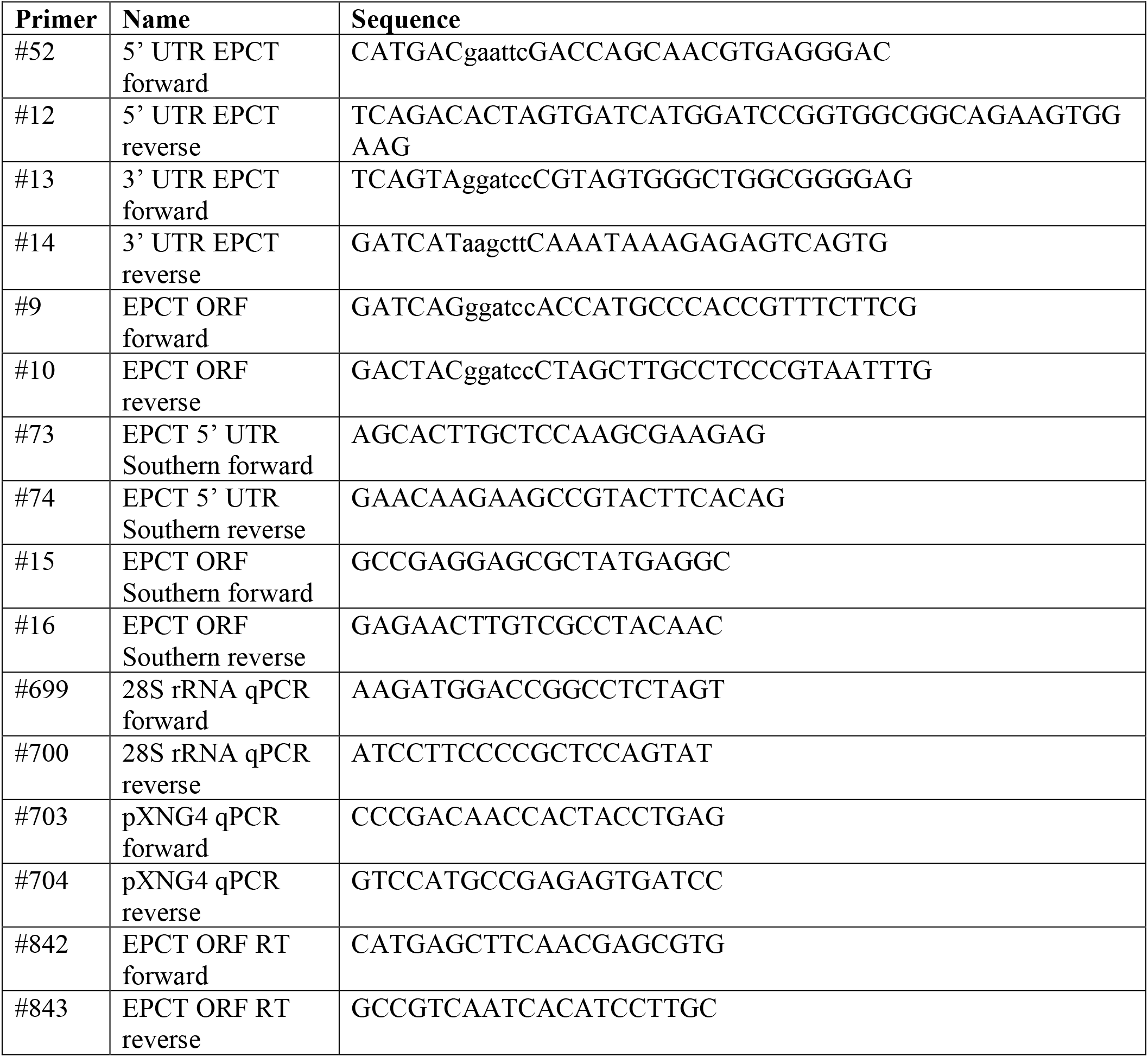
List of oligonucleotides used in this study. Sequences in lowercase represent restriction enzyme sites.

